# REC protein family expansion by the emergence of a new signaling pathway

**DOI:** 10.1101/2020.09.30.321588

**Authors:** Megan E. Garber, Vered Frank, Alexey E. Kazakov, Matthew R. Incha, Alberto A. Nava, Hanqiao Zhang, Jay D. Keasling, Lara Rajeev, Aindrila Mukhopadhyay

## Abstract

This report presents multi-genomes and experimental evidence that REC protein family expansion occurs when the emergence of new pathways give rise to functional discordance. Specificity between REC-domain containing response regulators with paired histidine kinases are under negative purifying selection, constrained by the presence of other bacterial two-component systems signaling cascades that share sequence and structural identity. Presuming that the two-component systems can evolve by neutral drift when these constraints are relaxed, how might the REC protein family expand when constraints remain intact? Using an unsupervised machine learning approach to observe the sequence landscape of REC domains across long phylogenetic distances, we find that within-gene-recombination, a subcategory of gene conversion, switched the effector domain, and consequently the regulatory context of a duplicated response regulator from transcriptional regulation by σ54 to σ70. We determined that the recombined response regulator diverged from its parent by positive episodic diversifying selection, giving rise to two new residues. Functional experiments of the parent of recombined response regulators in our model system, *Pseudomonas putida* KT2440, revealed that the parent and recombined response regulators sense and respond to carboxylic acids and that the two new residues in the recombined regulator form a new interaction interface and prevent crosstalk. Overall, our study finds genetic perturbations can create conditions of functional discordance, whereby the REC protein family can evolve by positive diversifying selection.

## Introduction

Large protein families are a group of proteins that share a common ancestor, showing strong sequence conservation across long phylogenetic distances (Heger and Holm 2003; Buljan and Bateman 2009; Finn et al. 2017; El-Gebali et al. 2019). Amino acid consensus motifs are used to understand protein family function and specificity; active-site residues are fixed and tell us about the chemistry the protein family performs or undergoes, while variable residues determine the protein’s specificity for different substrates or ligands. These variable, specificity-determining, residues form the molecular basis of tightly regulated signal transduction (Pao and Saier 1995; Capra and Laub 2012; Hirakawa et al. 2020), such as those found in phosphorelay receiver (REC) domains of response regulators, allowing for specific, context dependent, protein-protein interactions.

Members of the REC protein family (PF00072), function as phosphorelay receivers in bacterial signaling cascades called two-component systems. Most bacterial genomes encode dozens to over a hundred of these systems (Galperin 2006; Galperin 2018; Gumerov et al. 2020). Two component systems modulate important functions in bacterial pathogenesis, antibiotic resistance, nutrient use (e.g. carbon, nitrogen, or phosphate utilization), fitness in a microbiome context and a vast number of other functions that are responsive to signals (Gumerov et al. 2020; Hirakawa et al. 2020), making them attractive targets for addressing infection (Hirakawa et al. 2020), pathogenesis (Tiwari et al. 2017; Wang et al. 2021) and for beneficial applications in synbio and biotechnology (McClune et al. 2019; Schmidl et al. 2019; Volke et al. 2020). In the case of signaling systems and the REC protein family, despite large numbers of highly analogous systems encoded in a single genome, in these signaling cascades, membrane bound histidine kinases sense a signal, autophosphorylate, and with precise biochemical specificity, phosphorylate its cognate response regulator (Stock et al. 2000; Laub and Goulian 2007; Papon and Stock 2019). The phosphorylated cognate response regulator then regulates cellular functions, mainly via transcription, chemotaxis, or modulation of second messengers (Figure 1A). In response regulators, C-terminal REC domains are fused to a broad range of N-terminal effector domains, and predominantly function as transcription regulators (Figure 1B). Previous structural and biochemical studies of the REC protein family showed that variable amino acid sequences in active-site adjacent alpha-helices that from the interaction interface determine specificity with cognate histidine kinases to mediate highly specific, context dependent, phosphotransfer to the active-site aspartate residue (Varughese et al. 1998; Stock et al. 2000; Casino et al. 2009; Papon and Stock 2019).

**Figure 1:**
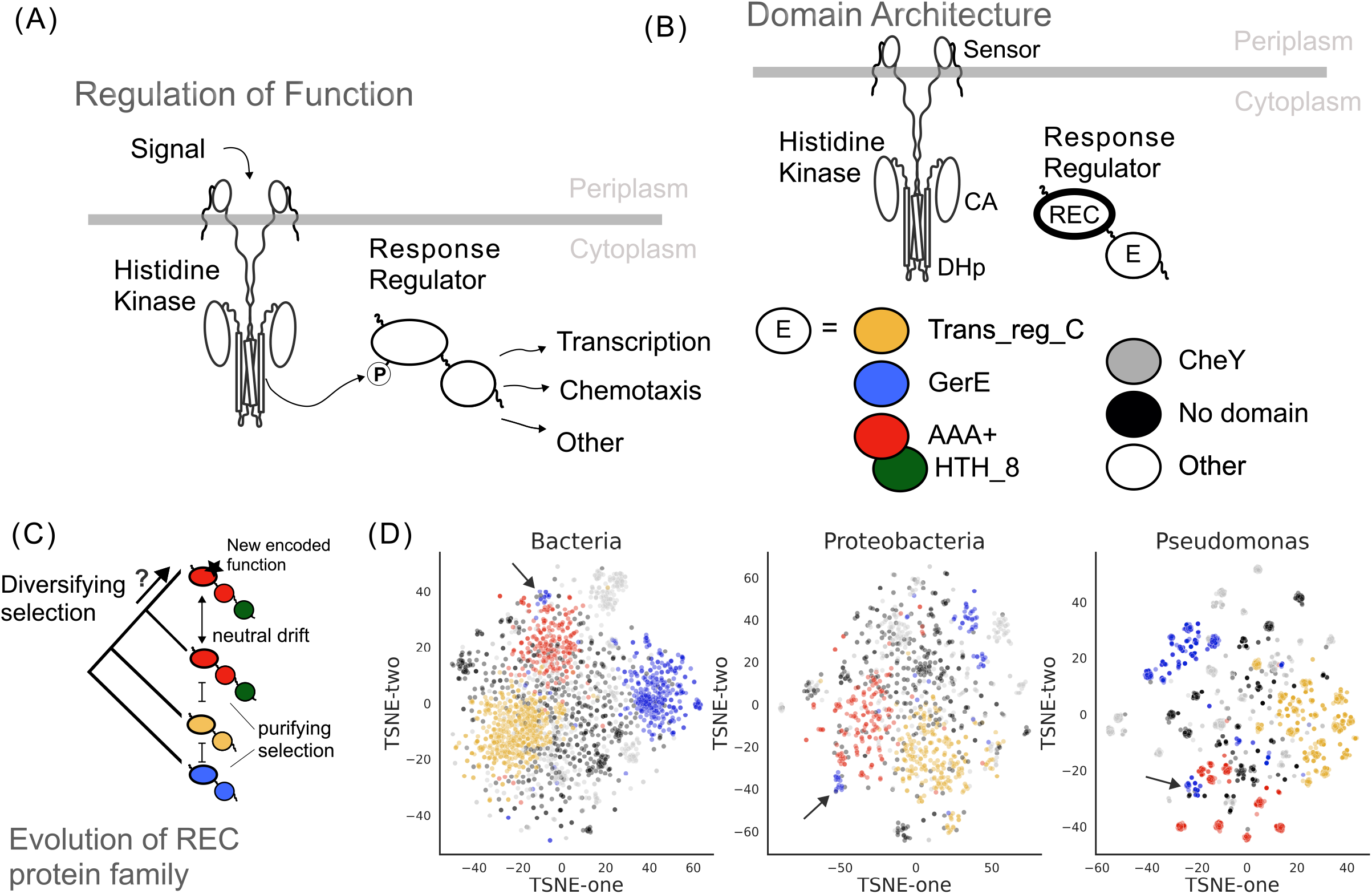
Functional context for insight into the evolution of the REC protein family. (A) REC protein family in the context of bacterial two-component systems: Membrane bound histidine kinases sense a signal, autophosphorylate, and phosphorylate, with precise biochemical specificity, a cognate response regulator. The phosphorylated cognate response regulator goes on to regulate cellular functions, including transcription, chemotaxis, or other functions. (B) Domain architecture of the two-component system (shown in A). The histidine kinases’ sensor, dimerization and histidine phosphotransfer (DHp), and catalytic (CA) domains coordinate the initial autophosphorylation in the two-component signaling cascade. The signal is transmitted by phosphotransfer to the cognate response regulators’ receiver (REC) domain (bolded oval). The effector domain (oval labeled “E”), completes the signaling cascade. Effector domains “E” are mostly DNA binding and the most abundant are Trans_reg_C (yellow), GerE (blue), AAA+ (red), and HTH_8 (green). Because HTH_8 domains often co-occur with AAA+ they are shown as overlapping red and green ovals - for simplicity, AAA+-HTH_8 domains are called AAA+ domains throughout the text. Effector domains with non-DNA binding function are shown as CheY (gray), other (white), and no domains (black). The effector domain color coding shown in this figure is used for all subsequent figures. (C) Model of the evolution of the REC protein family shown as a tree. We assume all REC domains share a common ancestor and that REC domains fused to the same effector domains are more closely related to one another than REC domains fused to different effector domains, showing regulators with distinct domain architectures and colors (refer to 1B) at the tips of the tree with coalescent branches. We know that genetic drift has occurred in the evolutionary history of the REC protein family when evolutionary constraints are relaxed, however, we do not know what causes REC domains to sample new combinations of sequences that encode new specificities when these constraints are intact. (D) Output from the dimensional reduction algorithm assesses the similarity between members of the REC protein family in Bacteria (leftmost panel), Proteobacteria (middle panel), and Pseudomonas (right panel) species, showing the relationship between REC domain sequence alignments with gap replacement and explaining 95% of the variation between the sequences. The REC domains were sampled from species found within the taxonomic rank (kingdom - Bacteria, phylum-Proteobacteria, or genera-Pseudomonas) labeled at the top of each plot. To control for overrepresentation of subclades due to more representative species with full genome sequences in any given subclade, species were sampled evenly from the taxonomic rank below the one represented in the plot title (e.g. The taxonomic rank below kingdom is phylum; the maximum of species from each bacterial phylum were randomly sampled to generate the bacteria dataset (see Table S1)). After species sampling, we identified all proteins with REC domains in each of the sampled species’ genomes and aligned their sequences. To read the plot, each point represents the sequence of a REC domain from the sampled genomes, points that are close together share sequence identity with each other. Each REC domain is annotated by the identity of its effector domain Trans_reg_C (yellow), GerE (blue), AAA+ (red), CheY (gray), other (gray outline), and no domains (black). Note that the location of REC domains is not fixed in the plot between independent runs of the algorithm, however, the relative distribution between the points is visually consistent between runs. The within-gene-recombination that changed parent REC domain fused to AAA+ domain to REC domain fused to GerE domain is apparent by the presence of two distinct blue clusters: the GerE cluster (blue) found close to the AAA+ cluster (red) resulted from within-gene-recombination of effector domains and is indicated by a black arrow.

How the REC protein family evolves is of interest due to their ubiquitous role in cellular signaling and regulation of cellular function. In the evolution of bacterial two-component systems, the interactions of cognate histidine kinases and response regulators are under negative purifying selection, constrained by functional overlap between signaling systems that share sequence identity at interaction interfaces (Siryaporn and Goulian 2008; Capra and Laub 2012; Capra et al. 2012; McClune and Laub 2020). Presumably, loss of a single two-component system, due to gene-loss or growth in conditions that suppress two-component system activity, relaxes these constraints, such that new two-component systems can sample the unused combinations of amino acids sequences that code for the interaction interfaces (Capra et al. 2012). It has been shown that while two-components sample new sequences, the phosphorelay function between cognate kinase and regulators do not break, and phosphotransfer levels can be maintained during the evolutionary process (Capra et al. 2010). This implies that REC protein family expansion occurs by neutral drift (Ohta 2001) when evolutionary constraints are relaxed, but we still do not understand if new combinations of sequences forming new interaction specificities are sampled when these constraints remain intact (Figure 1C).

One challenge in understanding the evolution of large protein families is parsing relationships between members of the protein family from amino acid sequences alone. Previous studies (Pao and Saier 1995; Chen et al. 2004; Ashby and Houmard 2006) using phylogenetic and clustering methods to parse molecular information from REC domains from species that span short evolutionary distances (population of a species or species within a genera) showed that members of the REC protein family resulted from independent gene-duplications, sharing a common ancestor in bacterial history. Studies comparing domain architectures of REC domains from extant species (Galperin 2005; Galperin 2010; Galperin 2018) suggest that REC domains evolve vertically, passing the fused effector domains to daughter REC domains (Figure 1C). To overcome the challenge in understanding how REC protein family expansion occurs over long phylogenetic distances, we build upon insights from these previous studies, making use of the over 200,000 sequenced bacterial genomes, to develop a method that can parse the evolution of the sequenced REC domains in extant, sequenced species (Gumerov et al. 2020).

In this study, we ask whether neutral genetic drift alone or another evolutionary mechanism can explain how new members of protein families emerge. Using an unsupervised machine learning strategy, we explore the amino acid sequence landscape of the REC protein family. We identify a subfamily of the REC protein family that emerged and expanded after a within-gene-recombination event that changed the effector domain of this REC subfamily. We show that functional discordance between parent and daughter signaling pathways, which occurred after the within-gene-recombination event, gave rise to a new structural interface that both facilitates phosphotransfer and prevents crosstalk.

## Results

To understand how large protein families expand and evolve, we applied an unsupervised machine learning (ML) algorithm that uses dimensional reduction to group similar domains based entirely on the identities and positions of amino acids sequences in an alignment. We established this strategy for large protein families, using the REC protein family (PF00072). To ensure diverse sampling of REC domains across the bacterial clade, we randomly sampled bacterial genomes from three taxonomic ranks, kingdom (Bacteria), phylum (Proteobacteria) and genera (*Pseudomonas*), generating independent datasets of REC sequences. To control for misgrouping due to gaps in the sequence alignments, we applied the algorithm to REC sequence alignments with and without gap replacement by randomly generated amino acids. Using t-Distributed Stochastic Neighbor Embedding algorithm(van der Maaten and Hinton 2008) with N principal components to explain 95% of the variation in the sequences, we projected the dimensionally reduced output of the ML algorithm in two-dimensions (Figures 1D, 2, Supplemental Figures 1-3) and color coded each REC domain (single point on the plot) with independent information about the effector domain it is fused to in nature. The data structure of the REC protein family becomes evident by highlighting the predominant effector domains that have transcriptional function, GerE, Trans_reg_C and AAA+, noting that few differences in data structure were apparent between REC sequences with and without gap replacement (Figures 1D, Supplemental Figure 1A). When we applied the ML-algorithm to scrambled amino acid sequence alignments with gap replacement, the REC domains appeared randomly distributed; however, REC domains with scrambled sequences without gap replacement appeared to be arranged non-randomly (Supplemental Figure 2). Determining that sequence ambiguity assigned by the hidden markov model alignment strategy (Zhang and Wood 2003) can result in more ambiguous alignments for REC amino acid sequences that share less sequence identity with the response regulators used to build the REC consensus model (Pao and Saier 1995), we decided that sequences with gap replacement would provide a data structure that was independent of the biases introduced by the alignment strategy. To demonstrate consistency of the method, we repeated the random sampling of REC sequences two more times and applied the strategy with gap replacement to each sampling, each showing similar data structures to the first sampling (Supplemental Figure 1 B,C). Together these results validate this strategy, allowing us to infer from it the evolutionary history of the REC protein family.

The REC protein family is divided into subfamilies, which we will call subclusters; each REC sequence is represented by a point in the two-dimensional sequence landscape (Figures 1D, 2, Supplemental Figures 1-3), and shares sequence identity with neighboring points. Subclusters that are closer together tend to share the identity of their fused effector domains (Figures 1B,C,D), indicating that the REC protein family evolved vertically, carrying effector domain architectures through to the next generation (Figure 1C). The separation between subclusters becomes more pronounced as we sample domains from lower taxonomic ranks (kingdom, to phylum, to genera), suggesting that the domains are under negative purifying selection. Based on the number of observed subclusters we can posit that despite evolutionary constraints, the REC protein family has expanded over the course of bacterial evolution, but we do not know whether protein families sample new combinations of amino acid sequences when these constraints remain intact. We hypothesized that crosstalk (overlap in specificity) between signaling pathways does not completely explain REC protein family expansion, because crosstalk is fairly common and does not always result in loss in fitness. For diversifying selection to occur, the signaling system would need to cause adverse effects on cellular function, such as those that can result from within-gene-recombination (Ohta 1991; Björklund et al. 2005; Forslund et al. 2008; Chan, Darling, et al. 2009; Chan, Beiko, et al. 2009; Forslund et al. 2019) (a subcategory of gene conversion, also known as domain swapping or domain shuffling).

Within-gene-recombination has occurred many times throughout the evolution of the REC protein family, and has been previously documented (Pao and Saier 1995; Stephenson and Hoch 2002; Chen et al. 2004; Qian et al. 2008). We chose to focus on the largest, most pronounced instance of with-in-gene recombination in our dataset. This event can be visualized in the two-dimensional sequence landscape by the presence of two major clusters of REC sequences with GerE effector domains (colored in blue); the GerE cluster that lies near REC sequences with AAA+ effectors (colored in red) is the product of a within-gene-recombination event that changed the effector domain of a parent response regulator with a AAA+ effector to GerE (Pao and Saier 1995; Chen et al. 2004). We focus on this event for two reasons: (1) it is a clear example of protein family expansion, given that the two distinct clusters of GerE fused response regulators are highly visible in the two-dimensional sequence landscape; and (2) the recombination event is by default linked to functional discordance, because AAA+ and GerE domains regulate transcription with distinct sigma (σ) factors, σ70 and σ54, which are active in incompatible cellular contexts (Cases et al. 2003; Potvin et al. 2008; Ronneau and Hallez 2019; Casas-Pastor et al. 2021).

To determine whether within-gene-recombination can lead to protein family expansion, we needed to determine whether these proteins were under diversifying selection as a result of the within-gene-recombination event. Using our unsupervised ML strategy, we were able to find that the recombined response regulator likely emerged sometime in the proteobacteria lineage, based on the presence of a second GerE subcluster in proteobacteria and its absence in other phyla (Supplemental Figure 3A-B). More specifically, the GerE subcluster expanded in the subclades of alpha, beta, and gammaproteobacteria (Figure 2, Supplemental Figure 3), but were not observable in deltaproteobacteria. In the species tree-of-life (Hug et al. 2016), alphaproteobacteria predates beta and gammaproteobacteria; we therefore reasoned that the within-gene-recombination event had occurred sometime in the alphaproteobacteria lineage. This insight enabled us repurpose the REC sequence landscapes to home in on the REC protein family in alphaproteobacteria (Figure 2, Supplemental Figure 3C) and search for the REC sequences of interest (Figure 3A) for detection of differential rates of evolution (Figure 3B, Supplemental Figure 4). We found that the within-gene-recombination event caused rapid (or episodic) diversifying selection on the branch of response regulators that have GerE domains in extant species (Supplemental Figure 4A). The consensus between the REC sequences of the parent and recombined response regulators highlights the shared sequence identity between the parent REC domains fused to AAA+ domains and recombined REC domains fused to GerE domains, apart from two highly conserved residues, a leucine and a cysteine, in the recombined REC domains fused to GerE domains (Figure 3C-D). We were also able to independently verify that these two residues in the recombined REC domains fused to GerE domains were under positive selection, using a method that identifies individual sites subject to episodic diversifying selection (Murrell et al. 2012) (Supplemental Data 1). These results show that after the within-gene-recombination, rapid selection occurred in the recombined regulators, however it does not explain the roles of the leucine and cysteine residues in their functional context of mediating phosphotransfer from their cognate kinases.

**Figure 2:**
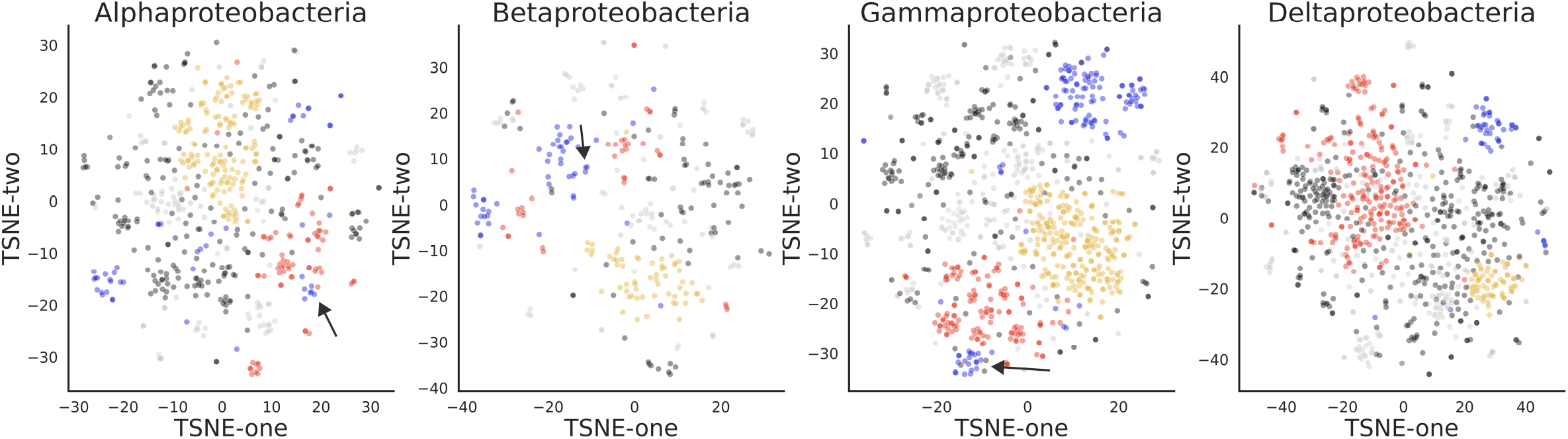
Within-gene-recombination that changed parent REC domain fused to AAA+ domain to REC domain fused to GerE domain occurred during the Alphaproteobacteria lineage. Output from the machine learning algorithm assesses the similarity between members of the REC protein family in (from left to right) Alphaproteobacteria, Betaproteobacteria, Gammaproteobacteria, Deltaproteobacteria species, showing the relationship between REC domain sequence alignments with gap replacement and explaining 95% of the variation between the sequences. The REC domains were sampled from species found within the taxonomic rank (class) labeled at the top of each plot. To control for overrepresentation of subclades due to more representative species with full genome sequences in any given subclade, species were sampled evenly from the taxonomic rank below the one represented in the plot title (e.g. the taxonomic rank below class is order; the maximum of species from each bacterial phylum were randomly sampled to generate the bacteria dataset (see Table S1)). After species sampling, we identified all proteins with REC domains in each of the sampled species’ genomes and aligned their sequences. To read the plot, each point represents the sequence of a REC domain from the sampled genomes, points that are close together share sequence identity with each other. Each REC domain is annotated by the identity of its effector domain Trans_reg_C (yellow), GerE (blue), AAA+ (red), CheY (gray), other (gray outline), and no domains (black). Note that the location of REC domains is not fixed in the plot between independent runs of the algorithm, however, the relative distribution between the points is visually consistent between runs. The within-gene-recombination that changed parent REC domain fused to AAA+ domain to REC domain fused to GerE domain is apparent by the presence of two distinct blue clusters: the GerE cluster (blue) found close to the AAA+ cluster (red) resulted from within-gene-recombination of effector domains and is indicated by a black arrow.

**Figure 3:**
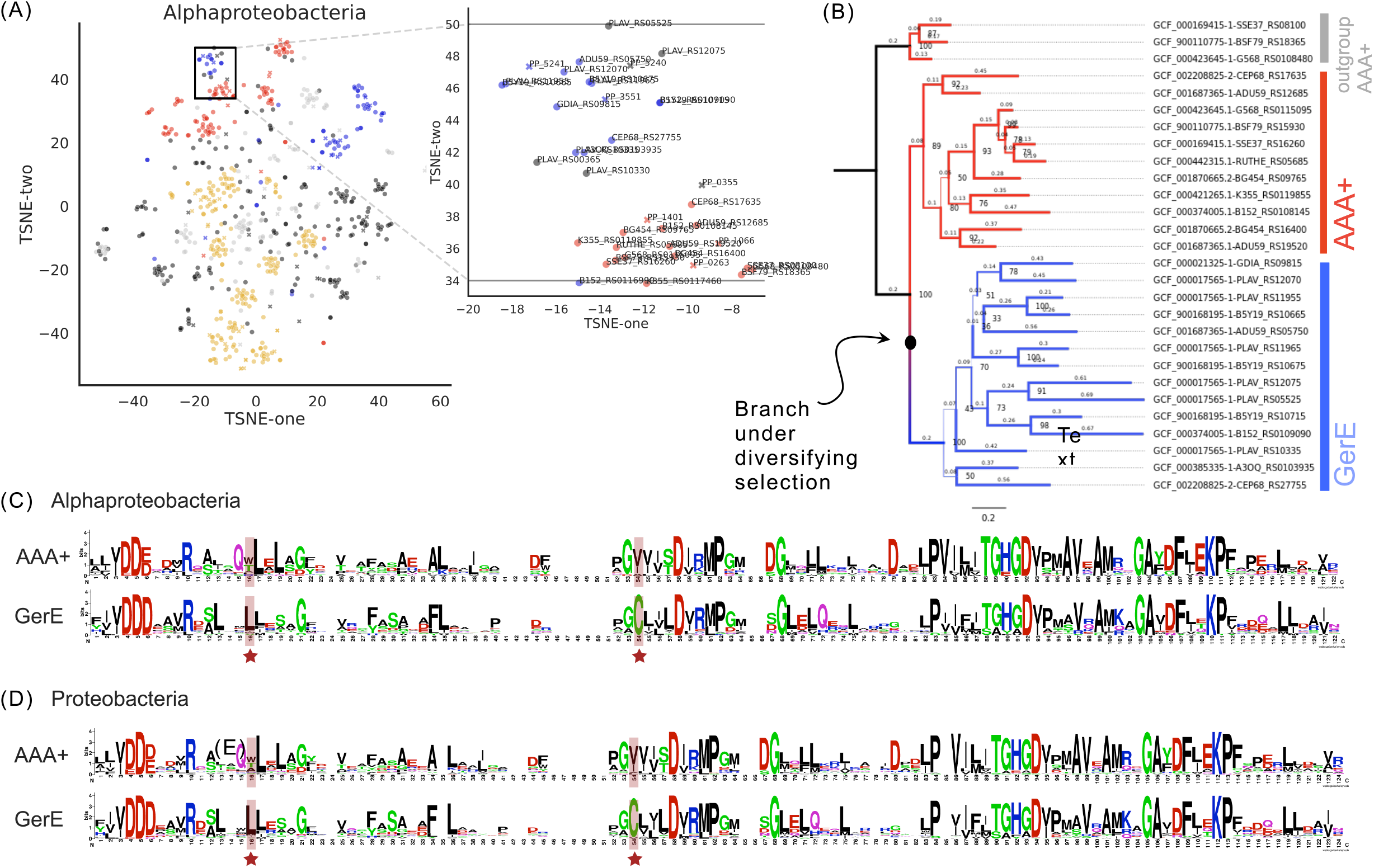
Episodic diversifying selection in the REC domain occurred after within-gene-recombination that changed parent REC domain fused to AAA+ domain to REC domain fused to GerE domain. (A) Output from the machine learning algorithm assesses the similarity between members of the REC protein family in Alphaproteobacteria species (circles) and *Pseudomonas putida* KT2440 (x’s). The sampled Alphaproteobacteria species are the same as those in Figure 2 (left-most panel); note that the location of REC domains is not fixed between independent runs of the algorithm, however, the relative distribution between the points is consistent between runs. The within-gene-recombination, which is apparent in Alphaproteobacteria by the presence of two distinct blue clusters: the GerE cluster (blue) found close to the AAA+ cluster (red) resulted from within-gene-recombination of effector domains, is indicated by a black box. The zoom panel of the black box shows the protein names of the REC domains within the box that are closely related to each other and have undergone a within-gene-recombination that switched effector domains. (B) The REC domains within the black box (Figure 3A) from alphaproteobacteria species (circles) are used to construct a domain tree. The DNA sequences of the REC domains were aligned with MAFFT. Using GCF_000169415-SSE37_RS08100 as an outgroup, we built the representative domain tree with iqtree, and annotated the branch colors red if the domain’s effector identity was AAA+ or blue if the domain’s effector identity was GerE (or no domain). Bootstrap supports are labeled at the nodes and shown as branch thickness. We independently determined that the recombined REC domains with GerE (or no domain) effectors were under episodic diversifying selection. The domain tree also shows that the parent REC domains with AAA+ effectors (red) independently duplicated and diverged after the within-gene-recombination. (C, D) The strategy applied to find the REC domains within the box (Figure 3A) was applied to a find parent REC domains with AAA+ effectors and recombined REC domains with GerE (or no domain) effectors for more Alphaproteobacteria species (C) and Proteobacteria species (D). The amino acid sequences of the REC domains were aligned with MAFFT and separated in two distinct groups (AAA+ or GerE) based on the identity of their effector domains. The alignments of each group were used to generate WebLogo consensus motifs revealing two residues, leucine and cysteine (in this alignment, L16 and C54 - highlighted with red background and red star below), are conserved in the GerE groups. Sequence alignments of AAA+ groups show the parent REC domains differ at alignment residue 16 having either alanine, tryptophan, or threonine at that position

To understand how the leucine and cysteine residues shaped the expansion of the recombined response regulators, we needed to probe their molecular roles in interaction and specificity with their cognate kinases and their parent’s cognate kinsases. However, the function of these regulators, the input signals they respond to and the genes they regulate, were not known before this study. We set out to determine those functions, by pursuing functional experiments in *Pseudomonas putida* KT2440, which proved advantageous for several reasons (1) *Pseudomonas putida* is a model organism in bioremediation and synthetic biology applications (Loeschcke and Thies 2015; Jumper et al. 2021; Sharma et al. 2022); therefore, many genetics and functional genomics experiments have been applied to understand this strain’s broad metabolism, allowing us to rapidly phenotype regulators to determine their functions; (2) *P. putida* KT2440 is a gammaproteobacteria, and is among the species where we have identified the parent AAA+ response regulators and the recombined GerE response regulator (note that these regulators are not present in *Escherichia coli*). Two functional genomics screens were used to determine, (1) the media conditions that cause randomly barcoded transposon insertions (RB-Tn) mutants at positions of the response regulators to more grow slowly in a population of other RB-Tn mutants (Supplemental Figure 5A); and (2) the genomic location of the DNA binding sites the response regulators use to regulate transcription (Supplemental Figure 5B and supplemental description). RB-Tn mutants of the response regulators showed growth phenotypes in defined media with carboxylic acids as the sole carbon source and DNA binding sites were found upstream of genes related to carboxylic acid assimilation; we therefore proposed that the regulators might respond to carboxylic acids to transcriptionally regulate a set of genes involved in carboxylic acid assimilation (Figure 4A). We validated these hypotheses with GFP reporter screens (Figure 4B, Supplemental Figure 6A), finding that the recombined regulator in *P. putida* responds to butyrate and regulates beta oxidation, while the parent regulator responds to glutamate and regulates amino acid assimilation. The overlap in chemical structure between carboxylic acids (Figure 4A) adds context of mis-signaling by chemical sensing to our proposed model for diversifying selection of the highly conserved residues in the recombined regulator, but we needed to directly investigate whether the evolved residues had a role in breaking interaction between the parent and recombined regulators’ signaling pathways.

**Figure 4:**
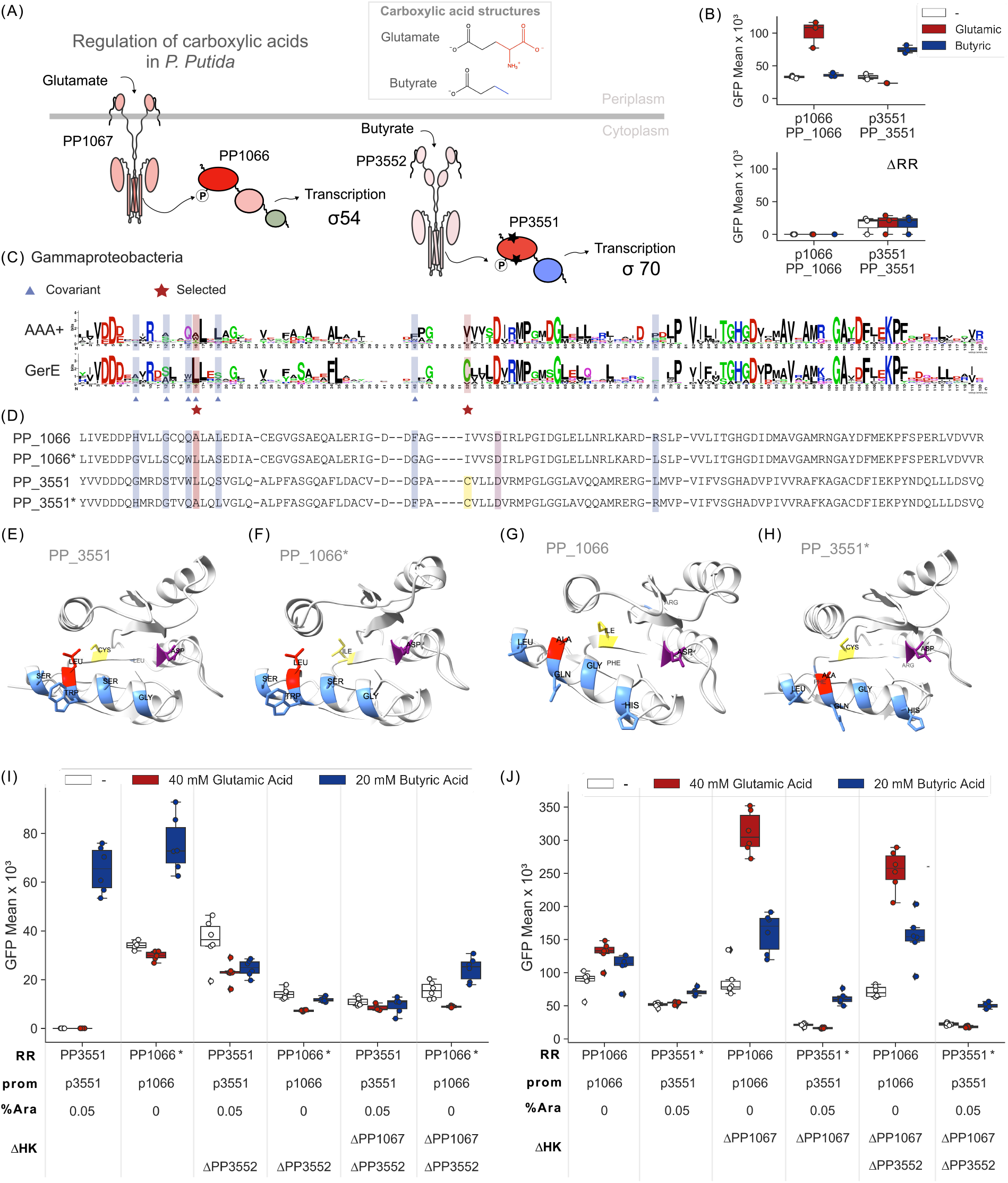
Parent and recombined response regulators in P. Putida KT2440 are regulators of carbon assimilation and evolved by episodic diversifying selection to break an overlap in specificity. (A) Model of regulation of carboxylic acids in *P. putida* KT2440 by the parent (PP1066) and recombined (PP3551) response regulators determined by functional genomics experiments.The parent two-component system (PP1067/PP1066), colored in red is activated by glutamate, and regulates genes for glutamate assimilation with AAA+ dependent σ54 transcription machinery. The recombined two-component system (PP3552/PP3551), also colored in red, is activated by butyrate, and regulates genes for beta oxidation with GerE dependent σ70 transcription machinery (the recombined GerE domain is colored blue). Based on the presence of transmembrane or sensing domains in the histidine kinases, we model PP1067 as membrane bound with a dCache domain for sensing glutamate, and PP3351 as cytosolic with two PAS domains for sensing butyrate. The residues in PP3551 REC domain under diversifying selection are represented by black stars on the PP3551 REC domain. Chemical similarity between glutamate and butyrate, which provides functional context to the proposed mechanism of functional discordance that caused diversifying selection, is shown in a box at the top right (B) *P. putida* KT2440 transformed with a GFP reporter plasmid for regulation by PP1066 or PP3551 were grown in defined glucose media with or without (white) glutamic acid (red) or butyric acid (blue). Top panel shows the reporter plasmid for PP1066 is upregulated when grown with glutamic acid and the reporter plasmid for PP3551 is upregulated when grown with butyric acid. Bottom panel shows reporters for PP1066 and PP3551 are respectively not activated if the regulators, PP1066 or PP3551 are deleted from the genome. Center line, median; box limits, upper and lower quartiles; whiskers, 1.5x interquartile range; points with black diamonds, outliers; n = 3. (C) The same strategy that was applied to find the REC domains within the black box (Figure 3A) was applied to find parent REC domains with AAA+ effectors and recombined REC domains with GerE (or no domain) effectors for Gammaproteobacteria species. The amino acid sequences of the REC domains were aligned with MAFFT and separated in two distinct groups (AAA+ or GerE) based on the identity of their effector domains. The alignments of each group were used to generate WebLogo consensus motifs, which revealed that two residues, leucine and cysteine (in this alignments, L16 and C52 - highlighted with red background and red star below), are conserved in the GerE groups. The 6 residues in the REC domain that tend to covary in Pseudomonas species are highlighted with blue background and blue triangle below, noting that L16 is both covariant and selected, while C52 is only selected. Sequence alignments here and in Figure 3C,D show the parent REC domains differ at alignment residue 16 having either alanine (PP1066: the parent regulator), tryptophan (PP1401), or threonine (PP0263) at that position. (D) Alignment of PP1066 and PP3551 REC domains. Covariant (blue), selected (yellow), covariant and selected (red), active aspartate (purple) residues are highlighted by the indicated background color. PP1066* and PP3551* match the sequence of PP1066 and PP3551, respectively, but are switched at the covariant positions (blue). These sequences test whether the specificity of PP1066 and PP3551 can be switched. (E,F,G,H) To show the structural impact of specificity switching, we modeled the structure of PP3551 (E), PP1066* (F), PP1066 (G), PP3551*(H) with the structural prediction of the full length protein using AlphaFold. Covariant (blue), selected (yellow), covariant and selected (red), active aspartate (purple) residues are highlighted. The covariant residues in PP3551 (E), including the selected leucine residue L16, is coded in PP1066* (F), the selected cysteine residue (C52) colored yellow is not. PP3551* is coded with specificity determining residues of PP1066, replacing selected leucine and covariant residue with alanine (L16A), but maintains its selected but not covariant cysteine residue (C52). (I) P. putida KT2440 ΔPP1066ΔPP3551 transformed with a GFP reporter plasmid for regulation by response regulators (RR) PP3551 and PP1066* were grown in defined glucose media with or without (white) glutamic acid (red) or butyric acid (blue). Percent arabinose (% ara) used to induce expression of heterologous response regulators and whether the respective cognate histidine kinases (ΔHK) are deleted from the genome is indicated below the plot. Center line, median; box limits, upper and lower quartiles; whiskers, 1.5x interquartile range; points with black diamonds, outliers; n = 6. (J) P. putida KT2440 ΔPP1066ΔPP3551 transformed with a GFP reporter plasmid for regulation by response regulators (RR) PP1066 and PP3551* were grown in defined glucose media with or without (white) glutamic acid (red) or butyric acid (blue). Percent arabinose (% ara) used to induce expression of heterologous response regulators and whether the respective cognate histidine kinases (ΔHK) are deleted from the genome is indicated below the plot. Center line, median; box limits, upper and lower quartiles; whiskers, 1.5x interquartile range; points with black diamonds, outliers; n = 6.

The domain tree of the parent and recombined regulators in alphaproteobacteria revealed that positive diversifying selection occurred after the emergence of the recombined regulator, but also revealed that the parent regulator underwent a second and third duplication event after the initial within-gene-recombination (Figure 3, Supplemental Figure 4). We did not detect instances of episodic diversifying selection on these post-recombination, duplicated REC domain branches, and thus determined they diverged from their parent REC domain by neutral genetic drift. From the functional genomics experiments, we determined that these regulators are also regulated by carboxylic acids and regulate carboxylic acid assimilation (Supplemental Figures 5,6A-B), raising an important distinction between these regulators that duplicated after the recombination event and the recombined regulators, despite the specificity overlap of sensing and responding to carboxylic acids with the parent, the post-recombination duplicated regulators diverged by genetic drift, they were not under negative purifying selection, nor positive diversifying selection. We believe this observation strengthens our hypothesis that specificity overlap and crosstalk is not disfavored, and that diversifying selection only occurs when the additional condition of functional discordance is met.

To understand if functional discordance following with-in-gene recombination plays a role in evolution in the REC protein family, we asked whether the identified mutations in the recombined regulator: (1) enables it to interact specifically with its cognate kinase; and (2) protects it from crosstalk with its parent regulator’s cognate kinase. To answer these questions, we applied a specificity-switching assay, in which we switched the six residues that tend to co-vary with cognate kinases across *Pseudomonads* (Supplemental Figure 7); the list of six residues included the conserved leucine residue, but excluded the conserved cysteine residue (Figure 4C-H). We found that the changes to the sequence of PP_1066, the parent regulator, were sufficient to switch its specificity to the recombined regulator’s cognate kinase (Figure 4I), however the same mutations were not sufficient to switch the specificity of PP_3551, the recombined regulator (Figure 4J). We hypothesized that the capability to switch by PP_1066, and inability to switch by PP_3551 was intrinsic to the role of the leucine and cysteine residues in regulating specificity and cros-talk.

To understand the physical role of leucine and cysteine in PP_3551 in connection to the observed trends in the specificity-switching experiment, we turned to the structural prediction of PP_3551, PP_1066, and their respective mutants determined by AlphaFold (Jumper et al. 2021). In PP_3551, the conserved leucine residue (Figure 4E-F, Supplemental Figure 8) sits inside of the alpha helix that canonically interacts directly with its cognate kinase. We can infer that when we switched PP_1066 with covarying residues from PP_3551, we encoded the new specificity, and enabled PP_1066 to interact with the cognate kinase of PP_3551 (Figure 4I), showing that the new interface is responsible for the interaction of PP_3551 with its cognate kinase. On the other hand, the cysteine residue lies within the same beta-sheet as the active aspartate residue that is canonically phosphorylated by a cognate kinase (Figure 4G-H). When we switched the residues of PP_3551 we left the cysteine residue intact, because it was not among the six covarying residues. The specificity switch PP_3551 mutant’s interaction site is almost identical to PP_1066 with the key difference being this cysteine residue. Because the specificity switched PP_3551 mutant did not interact with the cognate kinase of PP_1066 (Figure 4J), we reasoned that the cysteine residue may be responsible for preventing crosstalk with its parent, however further studies are need to reinforce this hypothesis. Therefore, the leucine residue plays a role in making contact with the cognate kinase of the recombined regulator, facilitating phosphotransfer, while we expect the cysteine residue plays a role in protecting the recombined regulator from interaction with the recombined regulator’s parent’s cognate kinase. Here we show that positive selection occurred after the within-gene-recombination event, forging new mechanisms of interaction specificity and protection against crosstalk.

## Discussion

The REC protein family undergoes three mechanisms of evolution, neutral genetic drift, negative purifying selection, and as determined by this study, positive diversifying selection (Figure 5A,B). Prior to this study, it was understood that neutral drift occurs when crosstalk between functionally overlapping signaling systems does not affect cellular fitness (Siryaporn and Goulian 2008; Groban et al. 2009; Capra et al. 2010; Siryaporn et al. 2010; Haushalter et al. 2015). It was also established that negative selection occurs when overlap in specificity between signaling systems results in dysregulation of cellular function affecting cellular fitness (Capra et al. 2012). This study demonstrates that REC protein family expansion can occur by positive diversifying selection, and has occurred in a test case, in which a new signaling system emerged by within-gene-recombination and functional overlap between the old and new signaling systems affected cellular fitness. Specifically, our results show that a within-gene-recombination in a duplicated response regulator of carbon assimilation changed the context in which this new regulator performed its transcriptional function (changing the effector domain from its parent regulator’s σ54 dependent transcription to σ70 dependent transcription). We show that specificity and functional overlap between the recombined response regulator and its parent gave rise to two new mutations in the recombined response regulator that form a new interface of interaction and prevents crosstalk with its parent response regulator (Figure 5A) by rapid diversifying selection.

**Figure 5:**
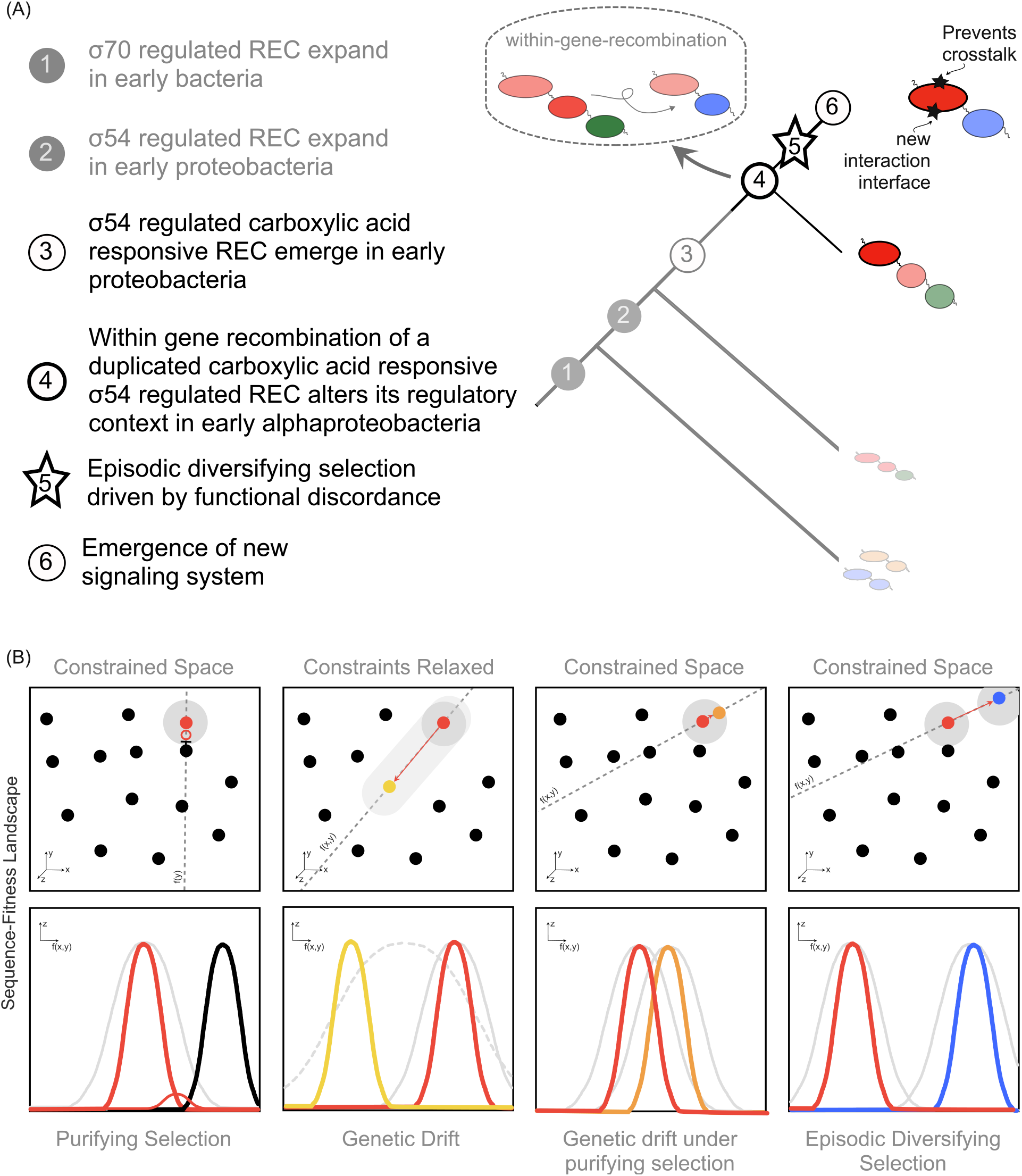
Models of the evolution of the REC protein family. (A) Model of the evolution of the REC protein family shown as a tree, updated by this study. We assume all REC domains share a common ancestor and that REC domains fused to the same effector domains evolved vertically, showing regulators with distinct domain architectures and colors (refer to 1B) at the tips of the tree with coalescent branches. Note that the order of events in steps 1-3 are uncertain. We model the findings of this study that diversifying selection and sequence divergence occurred after the within-gene-recombination that changed a parent REC domain fused to AAA+ domain to a recombined REC domain fused to a GerE domain. Because we determined that the parent and recombined REC domains regulate similar cellular responses and the recombined REC domains activate transcription under a new sigma factor regulatory contexts, we propose functional discordance gave rise to episodic diversifying selection, changing the interaction specificity and preventing the recombined REC domain from activating unwanted cellular responses that result from crosstalk. Changes that have broad cellular consequences, specifically those that cause functional discordance between members of large protein families can give rise to diversifying selection and lead to protein family expansion. (B) Models of the evolution of the REC protein family in a sequence-fitness landscape. Top and bottom panels show the sequence-fitness landscape as projections of sequence and fitness. The top panes show the x,y, plane representing the available combinations of amino acid sequences to the REC protein family, where z is the fitness (or distribution of REC domains with amino acid sequences defined by the x,y plane). If the fitness of a domain is high, it is shown as a solid black point on the plot, the boundaries of the fitness (or possible combinations of amino acid sequences that allow the domain to maintain its specificity) for the domains that are colored in red, yellow, orange, or blue are shown as a gray radius. Pointed arrows show the direction of positive selection or neutral drift, blunted arrows show the direction of negative selection. The bottom panes show the fitness as a function of the cross-section in the x,y plane (f(x,y)) (dashed line in the top panel) of the domain that are colored red, yellow, orange, or blue. Each column shows a different mode of evolution discussed in this report (from left to right: purifying selection, genetic drift, genetic drift under purifying selection, and episodic diversifying selection). Purifying selection selects against new REC domains that interfere with the specificity and function of existing REC domains. Genetic drift occurs when constraints in the sequence landscape are relaxed due to loss or suppressed activity of other REC domains that would otherwise constrain REC protein family expansion. Genetic drift under purifying selection can be tolerated when overlap in sequence specificity is not disadvantageous for the species in which the new sequence arises, however, this study shows that episodic diversifying selection occurs when the overlap in sequence specificity leads to functional discordance.

In this study we applied an unsupervised machine learning strategy to study the protein sequence landscape of the REC protein family. One of the benefits of this strategy, over previously applied strategies(Pao and Saier 1995; Chen et al. 2004; Ashby and Houmard 2006; Forslund et al. 2008; Forslund et al. 2019), is that it enabled us to characterize the relationship between REC protein family members over long evolutionary distances. The alternative approach of inferring evolution of a large protein family across long phylogenetic distances from a domain tree typically results in long branch distances, and uncertainty of branch positions (Efron et al. 1996; Simon 2022). For this reason, it is challenging to build reliable domain trees for large protein families, specifically the REC protein family, which we estimate has between 10-100 million members in sequenced bacteria (Gumerov et al. 2020). We overcame this challenge by repurposing the ML strategy to find closely related REC domains over a defined evolutionary distance (Figure 3A), after which we could build a reliable domain tree (Figure 3B) and apply methods to detect differential rates of evolution (Supplementary Figure 4A). One pitfall of the ML strategy is that the points on resultant maps do not maintain their positions for the same datasets between runs, however the relative position of the points for the same datasets are maintained between runs (van der Maaten and Hinton 2008)). Further development of this strategy, specifically implementing a bootstrapping method, will enhance the overall reliability in its application to the REC and other protein families. This tool will also be useful for distinguishing the relationships between orthologous and paralogous response regulators (or other large protein families) across species of interest.

Based on this study, we propose that diversifying selection in the REC protein family occurs when two conditions are met: (1) overlap in specificity between signaling cascades (for signals or interaction interfaces); and (2) the outcome of the regulation negatively impacts cellular fitness. Results from this study and others (Siryaporn and Goulian 2008; Groban et al. 2009; Capra et al. 2010; Noriega et al. 2010; Siryaporn et al. 2010; McClune et al. 2019) suggest that functional overlap between two-component systems that occur after duplication does not always result in negative purifying selection (Figure 5B). Specifically, in our study, we found that the parent response regulator duplicated and diverged a second and third time after the selection of the recombined regulator (Figure 3B, Supplemental Figure 4). Although these new regulators were presumably subjected to an inherent overlap in specificity and function with the parent regulator as they sampled sequences in their trajectory to new specificity, we determined they were not under positive diversifying selection, providing additional evidence that overlap in specificity can be tolerated by the cell (Figure 5B). The recombined regulator had comparable overlap in specificity with its parent regulator to these other duplicated response regulators - then why did the recombined regulators and regulators that duplicated and diverged after the recombination show different rates of evolution? After the within-gene-recombination changed the effector domain from its parent regulator’s σ54 dependent transcription to σ70 dependent transcription the recombined and parent response regulators regulated their overlapping function under the incompatible regulatory contexts of σ54 and σ70 (Cases et al. 2003; Potvin et al. 2008; Ronneau and Hallez 2019; Casas-Pastor et al. 2021). Although we cannot rule out the possibility that diversifying selection occurred for a different reason, the simplest explanation for the evolution of the recombined response regulator, is that differences in regulation by the sigma factors negatively impacted cellular fitness, and were the source of the functional discordance that gave rise to the selected mutations in the recombined regulator (Figure 5A).

This study establishes positive diversifying selection as a mechanism of the evolution for the REC protein family, however future studies are needed to determine the role of the selected cysteine residue in the recombined regulator. Our results show the conserved cysteine residue was positively selected (Figure 3), and experimental results allow us to speculate that this residue prevents interaction between the recombined regulator and its parent regulator, however, because we rely on the negative result that PP_3551 could not switch its specificity (Figure 4J), these experiments do not definitively confirm this hypothesis. Future studies are needed to confirm whether the conserved cysteine residue prevents crosstalk.

In our model for diversifying selection of a test case in the REC protein family, duplication followed by recombination led to an overlap in signals and outputs under competing regulatory contexts giving rise to residues that facilitate a new interaction and prevent crosstalk. We propose that diversifying selection is a mechanism of evolution for large protein family expansion if conditions of functional discordance are met, showing that within-gene-recombination can facilitate such conditions. In comparison to expansion in protein families where evolution is directly influenced by an immediate requirement; examples being dynamic environmental factors that need response in immune systems proteins (e.g. toxin-antitoxin (Jurėnas et al. 2022); Ig (Gaebler et al. 2021)) or small molecule sensing protein families (e.g. GPCRs/olfactory receptors (Bargmann 2006; Vandewege et al. 2016); efflux pumps (Henderson et al. 2021), the REC protein family is a midpoint in signaling cascades and immediate fitness or “arms race” factors may not be expected not influence its expansion. However, our study indicates that even for this class of large protein families positive selection occurs, not by changing environmental conditions - as we might have predicted - but instead due to genetic perturbations like recombination events. It should be noted that although recombination can be a source for functional discordance, as it is in this study’s test case, recombination can be neutral and will not always result in functional discordance. Further exploration of the REC sequence landscape or the sequence landscape of other protein families may elucidate whether other genetic perturbations create conditions for diversifying selection when evolutionary constraints remain intact. It will also be interesting to understand whether there are limitations on protein family expansion that occurs by neutral drift, even when evolutionary constraints are lifted. Overall, this study shows that positive diversifying selection is a mechanism in the expansion of the REC protein family and that it occurs by functional discordance. We believe this result is significant in understanding the evolution of the REC protein family, which directly modulates a vast range of cellular functions. This knowledge provides important context for assessing their role in microbial function, microbial communities and in engineering efforts in biosystems design and synthetic biology applications.

## Methods

### Two-dimensional visualization of protein family sequences

#### Sampling Genomes and creating REC domain databases

Databases of REC domains (PF00072) that were used to make all REC sequence landscapes were sampled from the microbial signal transduction database (MISTDB) (Gumerov et al. 2020) - a curated database of the signal transduction genes from over 200,000 bacterial genomes - using their API and custom functions written in python. To control for bias driven by genome availability for species of various taxonomic ranks (e.g gammaproteobacteria more represented than any other class in the proteobacteria phylum) we took a subset of the available data by randomly sampling species based on taxonomic rank (e.g. the taxonomic rank below kingdom is phylum; the maximum of species from each bacterial phylum were randomly sampled to generate the bacteria dataset (Table S1)). The following datasets were forced to include *P. putida* KT2440, which we used to track the parent and recombined regulator in the two-dimensional projections of the REC domains: Bacteria, Proteobacteria, *Pseudomonas*, and Alphaproteobacteria in Figure 3A. Again, using the MISTDB API, we used a custom function to generate a second database of all of the amino acid sequences of two-component system genes from the sampled species, while keeping a record of each protein’s metadata (e.g. fused domains). We then sampled this database to find all response regulators and their respective sequences to generate fasta files that include entries for each response regulator in the database.

#### REC domain alignments

We aligned the REC domain entries in the generated fasta files from the REC domain sampling functions using hmmalign (Zhang and Wood 2003; Finn et al. 2011) to the REC pfam consensus motif, PF00072 (Pao and Saier 1995). Using a custom python function, we linked the aligned domain back to its metadata in the REC domain database. When gap replacement was applied, gaps were replaced by a randomly generated amino acid. To generate scrambled datasets, we scrambled the individual sequences using a custom function that randomly samples residues and rejoins them in a new order while maintaining the original length of the alignment.

#### Projection of REC domain alignment variability in two-dimensional space

We then encoded the sequences into a discrete variable vector representation(Jing et al. 2020), by encoding each amino acid into a 21-dimensional zero vector (one dimension for each possible amino acid and a gap character), except for the index representing the amino acid, which is set to one. Each Lx21-dimensional matrix, where L is the length of the aligned protein sequence, is flattened to a Lx21-length vector. Each vector representing the alignment of a single protein sequence is stacked into a Nx(Lx21) matrix and dimensionally reduced to N dimensions to explain 95% of the variation by principal component analysis in the data before being reduced to Nx2 dimensions using the t-Distributed Stochastic Neighbor Embedding algorithm(van der Maaten and Hinton 2008). We approximated the perplexity, or the expected size of the clusters, as the number of clades in the taxonomic rank below the highest taxonomic rank queried (e.g. if we were looking at a kingdom rank, the next clade down is the phyla. The perplexity is then equal to the number of unique phyla in the dataset - see Table S2 (unique taxa in rank below).) Note that we found a binary encoding was sufficient for our use case, but another encoding may be more suitable for other large protein families.

Source code and data are available at https://github.com/mgarber21/Large_Protein_Families.git

### Isolating the parent and recombined regulators from randomly sampled REC domains in Alphaproteobacteria, Gammaproteobacteria, or Proteobacteria species

Using *P. putida* KT2440 to track the parent and recombined regulator in the two-dimensional projections of the REC domains in Alphaproteobacteria, Gammaproteobacteria, or Proteobacteria species, we defined boundaries in the projection and to isolate the candidate proteins. Using the DNA or amino acid sequences, we aligned the candidate domains (isolating the REC domain from the protein or protein coding sequence) using the MAFFT-LINSI algorithm from MAFFT v7.310(Katoh and Standley 2013). We then used IQTREE (Nguyen et al. 2015) to build the Alphaproteobacteria REC domain tree, using the aligned DNA sequences of the REC domains with a transversion model, empirical base frequencies, and discrete Gamma model (TVM+F+I+G4), also using outgroup species, GCF_000169415-1-SSE37_RS08100, and 1000 bootstraps. Using these same sequences, we applied an adaptive branch site rel test for episodic diversification, and detected individual sites subject to diversifying episodic selection using the online aBSREL (Kosakovsky Pond et al. 2011; Smith et al. 2015) and MEME (Murrell et al. 2012) tools from HyPhy in Datamonkey (Pond and Frost 2005; Delport et al. 2010; Weaver et al. 2018) with default parameters and according to the user instructions. Logs and results can be found in Supplemental Data 1. To generate amino acid consensus motif images for each subgroup, REC domains fused to either AAA+ or GerE, excluding outgroup AAA+ domains, we binned the aligned domains into their respective subgroups and generated a consensus motif using WebLogo (Crooks et al. 2004).

### Fitness experiments using a library of randomly barcoded transposon insertion (RB-Tn) mutants

As described in prior reports(Thompson et al. 2019), the *P. putida* KT2440 RB-TnSeq library, JBEI-1, was thawed, inoculated into 25 mL of LB medium supplemented with kanamycin, and grown to OD_600_ of 0.5. Three 1 mL samples were taken after this step to serve as t_0_ records of barcode abundance. The library was then washed via centrifugation and resuspension in an equal volume of MOPS minimal medium. The washed cells were then diluted 1:50 in MOPS minimal medium with 10 mM L-glutamate serving as the sole carbon source. The library was cultured in a 96-well deep well plate sealed with a gas permeable membrane (VWR, USA). The plate was shaken (700 rpm) in an INFORS HT Multitron (Infors USA Inc.) at 30 °C for 24 hrs. Duplicate 600 μL samples were then combined and BarSeq analysis was conducted as described previously(Wetmore et al. 2015; Rand et al. 2017; Incha et al. 2020). Single carbon source fitness data is available at http://fit.genomics.lbl.gov (Thompson et al. 2020).

### Automated DNA affinity purification with next generation sequencing (NGS)

#### DNA preparation for NGS

Pseudomonas isolates were cultured in either LB or minimal media (see Table S3 for strain specific minimal media recipes). Genomic DNA was purified with a promega wizard genomic preparation kit (Promega, Madison, WI). DNA was sheared with covaris miniTUBE (Covaris, Woburn, MA) to an average size of 200 bp. The DNA quality was confirmed by Bioanalyzer high sensitivity DNA kit (Agilent, Santa Clara). Sheared DNA was then adapter-ligated (AL) with NEBnext Ultra ii Library Preparation kit (New England Biolabs, Ipswich, MA). AL-DNA quality was again confirmed by Bioanalyzer high sensitivity DNA kit (Agilent, Santa Clara). AL-DNA was stored at −20 °C until required for downstream use.

#### Expression Strain Design

pet28 expression vectors with N-terminal 6x-His-tagged RRs were cloned by Gibson assembly(Gibson et al. 2009). Plasmid design was facilitated by j5 DNA assembly design(Hillson et al. 2012), see Table S4 for primers.

#### Automated DNA affinity purification

Quadruplicates of expression strains were grown in autoinduction media (Zyp-5052(Studier 2005)) at 37 °C, 250 RPM, for 5-6 hours and then transferred to grow at 17 °C, 250 RPM, overnight. Cell pellets were harvested and lysed at 37 °C for 1 hour in a lysis buffer (1X TBS, 100 μM PMSF (Millipore Sigma, Burlington MA), 2.5 units/mL Benzonase nuclease (Millipore Sigma, Burlington MA), 1 mg/mL Lysozyme (Millipore Sigma, Burlington MA)). Lysed cells were then clarified by centrifugation at 3214 x g and further filtered in 96-well filter plates by centrifugation at 1800 x g. To enable high-throughput processing, protein-DNA purification steps were performed with IMAC resin pipette tips (PhyNexus, San Jose, CA) using a custom automated platform with the Biomek FX liquid handler (Beckman Coulter, Indianapolis, IN). The expressed RRs were individually bound to metal affinity resin embedded within the IMAC resin pipette tips and washed in a wash buffer (1X TBS, 10 mM imidazole, 0.1% Tween 20). The bead bound RRs were then mixed with 60 μL of DNA binding buffer (1X TBS, 10 mM magnesium chloride, 0.4 ng/μL AL-DNA, with or without 50 mM acetyl phosphate (split into duplicates)). The protein bound to its target DNA was then enriched in an enrichment buffer (1X TBS, 10 mM imidazole, 0.1% Tween 20) and eluted in an elution buffer (1X TBS, 180 mM imidazole). The elution was stored at −20 °C for a minimum of one day and up to a week before proceeding to the NGS library generation. See supplementary methods for detailed protocol.

#### NGS Library Generation

3.2 μL of the elution from the previous step was added to 3.5 μL SYBR green ssoAdvanced (Biorad, Hercules, CA) and 0.15 μL of each dual indexed NGS primers. NGS libraries were prepared by following the protocols for fluorescent amplification of NGS libraries(Chiniquy et al. 2020). Pooled libraries were sequenced by Illumina NovaSeq 6000 SP (100 cycles) (Illumina, San Diego, CA).

#### DAP-seq data analysis

Sequenced reads were processed by a computational DAP-seq analysis pipeline as follows. Adapters and low-quality bases were trimmed and reads shorter than 30 bp were filtered out using Trimmomatic v.0.36(Bolger et al. 2014). The resulting reads were checked for contamination using FOCUS (Silva et al. 2014). Then the reads were aligned to the corresponding *Pseudomonas* spp. genome using Bowtie v1.1.2 (Langmead et al. 2009) with -m 1 parameter (report reads with single alignment only). Resulting SAM files were converted to BAM format and sorted using samtools v 0.1.19(Li et al. 2009). Peak calling was performed using SPP 1.16.0(Kharchenko et al. 2008) with false discovery rate threshold of 0.01 and MLE enrichment ratio threshold of 4.0. Enriched motifs were discovered in genome fragments corresponding to the peaks using MEME(Bailey et al. 2009) with parameters -mod anr -minw 12 -maxw 30 - revcomp -pal -nmotifs 1. Source code of the DAP-seq analysis pipeline is available at https://github.com/novichkov-lab/dap-seq-utils.

For conserved RRs with small numbers of high-confidence peaks (1-2 per genome), binding motifs were predicted manually by a comparative genomics approach. Orthologous RRs were identified by OrthoFinder2(Emms and Kelly 2019). For each of orthologous RRs, one genome fragment corresponding to the peak with the highest enrichment value was selected for motif search. Conserved motifs were discovered using the SignalX tool from the GenomeExplorer package(Mironov et al. 2000) with the “inverted repeat” option.

### Co-variance analysis

Cognate RRs and HKs from *Pseudomonas* and *E. coli* strains were identified as pairs if they were found neighboring each other in their respective genomes. DHp (HisKA), CA (HATPase_C), and REC (Response_reg) domain boundaries were determined with hmmsearch from HMMER v3.1b2(Mistry et al. 2013). Fasta files of concatenated DHp-CA-REC domains from cognate and randomized HK-RR pairs were aligned with the MAFFT-LINSI algorithm from MAFFT v7.310(Katoh and Standley 2013). Alignment files were then queried for coevolution with the ProDy *Evol* suite(Bakan et al. 2011; Bakan et al. 2014) in python and were plotted in a heatmap. The highest scoring residues > 1.1 were used to inform hypotheses for specificity switch strains.

### GFP reporter strain generation and assays

#### Knockout strain generation

1000bp homology fragments upstream and downstream of the target gene were cloned into plasmid pKS18. Plasmids were then transformed into *E. coli* S17 and then mated into *P. putida* via conjugation. Transconjugants were selected for on LB agar plates supplemented with 30 mg/ml kanamycin and 30 mg/ml chloramphenicol. Transconjugants were then grown overnight on LB medium and were then plated on LB agar with no NaCl that was supplemented with 10% (wt/vol) sucrose. Putative deletions were screened on LB agar with no NaCl supplemented with 10% (wt/vol) sucrose and LB agar plate with kanamycin. Colonies that grew in the presence of sucrose but had no resistance to kanamycin were further tested via PCR with primers flanking the target gene to confirm gene deletion.

#### GFP reporter strains

Promoter boundaries for p2453, p1400, p3553 were selected as the region just upstream of the gene’s start codon up until the start or stop codon of the next nearest gene. The promoters were cloned upstream of the gene encoding sfGFP on a broad host range plasmid with BBR1 origin and Kanamycin resistance, with Gibson cloning(Gibson et al. 2009). Primers in Table S4. The plasmids were transformed into *P. putida* KT2440 or *P. putida* KT2440 mutant strains by electroporation. Three biological replicates of each strain were cultured in LB and stored with 25% (vol/vol) glycerol at −80 °C. Complementation plasmids (GFP reporter plasmids with full length RR driven by pBAD promoter and constitutively expressed AraC) were combinatorially built leveraging Golden Gate cloning(Engler et al. 2008) and j5 DNA assembly design(Hillson et al. 2012) (diva.jbei.org), primers in Table S4. The plasmids were transformed into gene knockout strains of *P. putida* KT2440 by electroporation. 3-6 biological replicates of each strain were cultured in LB and stored with 25 % (v/v) glycerol at −80 °C.

#### RRs with switched specificity

Gene blocks (TWIST Biosciences, San Francisco, CA) of REC domains (Table S3) with co-varying mutations (co-variation score > 1.1 - see supplemental figure 7) were cloned into the complementation plasmids with Gibson assembly(Gibson et al. 2009). The plasmids were transformed into *P. putida* KT2440 or knockout strains of *P. putida* KT2440 by electroporation. Six biological replicates of each strain were cultured in LB and stored with 25 % (vol/vol) glycerol at −80 °C.

#### GFP reporter Assays

Reporter strains were adapted to M9 minimal media (MM) (see supplemental methods for strain specific minimal media recipes) supplemented with 0.5 % (wt/vol) glucose as the sole carbon source in 3 overnight passages, and stored in MM at −80 °C in 25 % (vol/vol) glycerol. Adapted strains were cultured in MM + 0.5 % (wt/vol) glucose and passaged to MM + 0.5 % (wt/vol) glucose with or without a second carbon-source (40 mM glutamic acid, 40 mM a-ketoglutaric acid, or 20 mM butyric acid, unless otherwise specified). Note that specificity switch strains have response regulators under pBAD promoters, which were first calibrated with 2-fold dose response of arabinose concentrations between 0-0.1% (Supplemental Figure 9). After 24-hours of growth, samples were diluted 1:100 in 1X PBS, and fluorescence was measured by flow cytometry on the BD Accuri C6 (BD Biosciences, San Jose, CA). Autofluorescence was gated out with FlowJo (BD Biosciences, San Jose, CA), using a non-fluorescent strain of *P. putida* KT2440 carrying an empty vector for reference (Supplemental Figure 10). To remove noise, the GFP mean for samples with less than 150 events after gating was set to 0. Otherwise, the GFP mean of the remaining events after gating was reported.

### AlphaFold predictions of wildtype and mutant response regulators

Full length amino acid sequences of wildtype PP_1066, and the respective specificity switching mutants of PP_3551 and PP_1066 were queried using ColabFold by AlphaFold (Jumper et al. 2021) using the default parameters.PDB files and prediction logs can be found Supplemental Data 2. The structural prediction of PP_3551 was retrieved from the AlphaFold deepmind database (Varadi et al. 2022). The pdb file from each protein’s highest ranking structure from AlphaFold was then visualized and annotated with Chimera X (Pettersen et al. 2021).

## References

Ashby MK, Houmard J. 2006. Cyanobacterial two-component proteins: structure, diversity, distribution, and evolution. Microbiol. Mol. Biol. Rev. 70:472–509.

Bailey TL, Boden M, Buske FA, Frith M, Grant CE, Clementi L, Ren J, Li WW, Noble WS. 2009. MEME SUITE: tools for motif discovery and searching. Nucleic Acids Res. 37:W202–8.

Bakan A, Dutta A, Mao W, Liu Y, Chennubhotla C, Lezon TR, Bahar I. 2014. Evol and ProDy for bridging protein sequence evolution and structural dynamics. Bioinformatics 30:2681–2683.

Bakan A, Meireles LM, Bahar I. 2011. ProDy: protein dynamics inferred from theory and experiments. Bioinformatics 27:1575–1577.

Bargmann CI. 2006. Comparative chemosensation from receptors to ecology. Nature 444:295–301.

Björklund AK, Ekman D, Light S, Frey-Skött J, Elofsson A. 2005. Domain rearrangements in protein evolution. J. Mol. Biol. 353:911–923.

Bolger AM, Lohse M, Usadel B. 2014. Trimmomatic: a flexible trimmer for Illumina sequence data. Bioinformatics 30:2114–2120.

Buljan M, Bateman A. 2009. The evolution of protein domain families. Biochem. Soc. Trans. 37:751–755.

Capra EJ, Laub MT. 2012. Evolution of two-component signal transduction systems. Annu. Rev. Microbiol. 66:325–347.

Capra EJ, Perchuk BS, Lubin EA, Ashenberg O, Skerker JM, Laub MT. 2010. Systematic dissection and trajectory-scanning mutagenesis of the molecular interface that ensures specificity of two-component signaling pathways. PLoS Genet. 6:e1001220.

Capra EJ, Perchuk BS, Skerker JM, Laub MT. 2012. Adaptive mutations that prevent crosstalk enable the expansion of paralogous signaling protein families. Cell 150:222–232.

Casas-Pastor D, Müller RR, Jaenicke S, Brinkrolf K, Becker A, Buttner MJ, Gross CA, Mascher T, Goesmann A, Fritz G. 2021. Expansion and re-classification of the extracytoplasmic function (ECF) σ factor family. Nucleic Acids Res. 49:986–1005.

Cases I, Ussery DW, De Lorenzo V. 2003. The σ^54^ regulon (sigmulon) ofPseudomonas putida. Environ. Microbiol. 5:1281–1293.

Casino P, Rubio V, Marina A. 2009. Structural insight into partner specificity and phosphoryl transfer in two-component signal transduction. Cell 139:325–336.

Chan CX, Beiko RG, Darling AE, Ragan MA. 2009. Lateral transfer of genes and gene fragments in prokaryotes. Genome Biol. Evol. 1:429–438.

Chan CX, Darling AE, Beiko RG, Ragan MA. 2009. Are protein domains modules of lateral genetic transfer? PLoS ONE 4:e4524.

Chen Y-T, Chang HY, Lu CL, Peng H-L. 2004. Evolutionary analysis of the two-component systems in Pseudomonas aeruginosa PAO1. J. Mol. Evol. 59:725–737.

Chiniquy J, Garber ME, Mukhopadhyay A, Hillson NJ. 2020. Fluorescent amplification for next generation sequencing (FA-NGS) library preparation. BMC Genomics 21:85.

Crooks GE, Hon G, Chandonia JM, Brenner SE. 2004. WebLogo: a sequence logo generator. Genome Res. 14:1188–1190.

Delport W, Poon AFY, Frost SDW, Kosakovsky Pond SL. 2010. Datamonkey 2010: a suite of phylogenetic analysis tools for evolutionary biology. Bioinformatics 26:2455–2457.

Efron B, Halloran E, Holmes S. 1996. Bootstrap confidence levels for phylogenetic trees. Proc Natl Acad Sci USA 93:13429–13434.

El-Gebali S, Mistry J, Bateman A, Eddy SR, Luciani A, Potter SC, Qureshi M, Richardson LJ, Salazar GA, Smart A, et al. 2019. The Pfam protein families database in 2019. Nucleic Acids Res. 47:D427–D432.

Emms DM, Kelly S. 2019. OrthoFinder: phylogenetic orthology inference for comparative genomics. Genome Biol. 20:238.

Engler C, Kandzia R, Marillonnet S. 2008. A one pot, one step, precision cloning method with high throughput capability. PLoS ONE 3:e3647.

Finn RD, Attwood TK, Babbitt PC, Bateman A, Bork P, Bridge AJ, Chang H-Y, Dosztányi Z, El-Gebali S, Fraser M, et al. 2017. InterPro in 2017-beyond protein family and domain annotations. Nucleic Acids Res. 45:D190–D199.

Finn RD, Clements J, Eddy SR. 2011. HMMER web server: interactive sequence similarity searching. Nucleic Acids Res. 39:W29–37.

Forslund K, Henricson A, Hollich V, Sonnhammer ELL. 2008. Domain tree-based analysis of protein architecture evolution. Mol. Biol. Evol. 25:254–264.

Forslund SK, Kaduk M, Sonnhammer ELL. 2019. Evolution of protein domain architectures. Methods Mol. Biol. 1910:469–504.

Gaebler C, Wang Z, Lorenzi JCC, Muecksch F, Finkin S, Tokuyama M, Cho A, Jankovic M, Schaefer-Babajew D, Oliveira TY, et al. 2021. Evolution of antibody immunity to SARS-CoV-2. Nature 591:639–644.

Galperin MY. 2005. A census of membrane-bound and intracellular signal transduction proteins in bacteria: bacterial IQ, extroverts and introverts. BMC Microbiol. 5:35.

Galperin MY. 2006. Structural classification of bacterial response regulators: diversity of output domains and domain combinations. J. Bacteriol. 188:4169–4182.

Galperin MY. 2010. Diversity of structure and function of response regulator output domains. Curr. Opin. Microbiol. 13:150–159.

Galperin MY. 2018. What bacteria want. Environ. Microbiol. 20:4221–4229.

Gibson DG, Young L, Chuang R-Y, Venter JC, Hutchison CA, Smith HO. 2009. Enzymatic assembly of DNA molecules up to several hundred kilobases. Nat. Methods 6:343–345.

Groban ES, Clarke EJ, Salis HM, Miller SM, Voigt CA. 2009. Kinetic buffering of cross talk between bacterial two-component sensors. J. Mol. Biol. 390:380–393.

Gumerov VM, Ortega DR, Adebali O, Ulrich LE, Zhulin IB. 2020. MiST 3.0: an updated microbial signal transduction database with an emphasis on chemosensory systems. Nucleic Acids Res. 48:D459–D464.

Haushalter RW, Groff D, Deutsch S, The L, Chavkin TA, Brunner SF, Katz L, Keasling JD. 2015. Development of an orthogonal fatty acid biosynthesis system in E. coli for oleochemical production. Metab. Eng. 30:1–6.

Heger A, Holm L. 2003. Exhaustive enumeration of protein domain families. J. Mol. Biol. 328:749–767.

Henderson PJF, Maher C, Elbourne LDH, Eijkelkamp BA, Paulsen IT, Hassan KA. 2021. Physiological functions of bacterial “multidrug” efflux pumps. Chem. Rev. 121:5417–5478.

Hillson NJ, Rosengarten RD, Keasling JD. 2012. j5 DNA assembly design automation software. ACS Synth. Biol. 1:14–21.

Hirakawa H, Kurushima J, Hashimoto Y, Tomita H. 2020. Progress Overview of Bacterial Two-Component Regulatory Systems as Potential Targets for Antimicrobial Chemotherapy. Antibiotics (Basel) 9.

Hug LA, Baker BJ, Anantharaman K, Brown CT, Probst AJ, Castelle CJ, Butterfield CN, Hernsdorf AW, Amano Y, Ise K, et al. 2016. A new view of the tree of life. Nat. Microbiol. 1:16048.

Incha MR, Thompson MG, Blake-Hedges JM, Liu Y, Pearson AN, Schmidt M, Gin JW, Petzold CJ, Deutschbauer AM, Keasling JD. 2020. Leveraging host metabolism for bisdemethoxycurcumin production in Pseudomonas putida. Metab. Eng. Commun. 10:e00119.

Jing X, Dong Q, Hong D, Lu R. 2020. Amino acid encoding methods for protein sequences: A comprehensive review and assessment. IEEE/ACM Trans Comput Biol Bioinform 17:1918–1931.

Jumper J, Evans R, Pritzel A, Green T, Figurnov M, Ronneberger O, Tunyasuvunakool K, Bates R, Žídek A, Potapenko A, et al. 2021. Highly accurate protein structure prediction with AlphaFold. Nature 596:583–589.

Jurėnas D, Fraikin N, Goormaghtigh F, Van Melderen L. 2022. Biology and evolution of bacterial toxin-antitoxin systems. Nat. Rev. Microbiol. 20:335–350.

Katoh K, Standley DM. 2013. MAFFT multiple sequence alignment software version 7: improvements in performance and usability. Mol. Biol. Evol. 30:772–780.

Kharchenko PV, Tolstorukov MY, Park PJ. 2008. Design and analysis of ChIP-seq experiments for DNA-binding proteins. Nat. Biotechnol. 26:1351–1359.

Kosakovsky Pond SL, Murrell B, Fourment M, Frost SDW, Delport W, Scheffler K. 2011. A random effects branch-site model for detecting episodic diversifying selection. Mol. Biol. Evol. 28:3033–3043.

Langmead B, Trapnell C, Pop M, Salzberg SL. 2009. Ultrafast and memory-efficient alignment of short DNA sequences to the human genome. Genome Biol. 10:R25.

Laub MT, Goulian M. 2007. Specificity in two-component signal transduction pathways. Annu. Rev. Genet. 41:121–145.

Li H, Handsaker B, Wysoker A, Fennell T, Ruan J, Homer N, Marth G, Abecasis G, Durbin R, 1000 Genome Project Data Processing Subgroup. 2009. The Sequence Alignment/Map format and SAMtools. Bioinformatics 25:2078–2079.

Loeschcke A, Thies S. 2015. Pseudomonas putida-a versatile host for the production of natural products. Appl. Microbiol. Biotechnol. 99:6197–6214.

van der Maaten L, Hinton G. 2008. Visualizing data using t-SNE. J Mach Learn Res 8:2579–2605.

McClune CJ, Alvarez-Buylla A, Voigt CA, Laub MT. 2019. Engineering orthogonal signalling pathways reveals the sparse occupancy of sequence space. Nature.

McClune CJ, Laub MT. 2020. Constraints on the expansion of paralogous protein families. Curr. Biol. 30:R460–R464.

Mironov AA, Vinokurova NP, Gelfand MS. 2000. Software for analysis of bacterial genomes. Mol Biol (NY) 34:222–231.

Mistry J, Finn RD, Eddy SR, Bateman A, Punta M. 2013. Challenges in homology search: HMMER3 and convergent evolution of coiled-coil regions. Nucleic Acids Res. 41:e121.

Murrell B, Wertheim JO, Moola S, Weighill T, Scheffler K, Kosakovsky Pond SL. 2012. Detecting individual sites subject to episodic diversifying selection. PLoS Genet. 8:e1002764.

Nguyen L-T, Schmidt HA, von Haeseler A, Minh BQ. 2015. IQ-TREE: a fast and effective stochastic algorithm for estimating maximum-likelihood phylogenies. Mol. Biol. Evol. 32:268–274.

Noriega CE, Lin H-Y, Chen L-L, Williams SB, Stewart V. 2010. Asymmetric cross-regulation between the nitrate-responsive NarX-NarL and NarQ-NarP two-component regulatory systems from Escherichia coli K-12. Mol. Microbiol. 75:394–412.

Ohta T. 1991. Role of diversifying selection and gene conversion in evolution of major histocompatibility complex loci. Proc Natl Acad Sci USA 88:6716–6720.

Ohta T. 2001. Molecular evolution: nearly neutral theory. In: John Wiley & Sons, Ltd, editor. eLS. Chichester, UK: John Wiley & Sons, Ltd.

Pao GM, Saier MH. 1995. Response regulators of bacterial signal transduction systems: selective domain shuffling during evolution. J. Mol. Evol. 40:136–154.

Papon N, Stock AM. 2019. Two-component systems. Curr. Biol. 29:R724–R725.

Pettersen EF, Goddard TD, Huang CC, Meng EC, Couch GS, Croll TI, Morris JH, Ferrin TE. 2021. UCSF ChimeraX: structure visualization for researchers, educators, and developers. Protein Sci. 30:70–82.

Pond SLK, Frost SDW. 2005. Datamonkey: rapid detection of selective pressure on individual sites of codon alignments. Bioinformatics 21:2531–2533.

Potvin E, Sanschagrin F, Levesque RC. 2008. Sigma factors in Pseudomonas aeruginosa. FEMS Microbiol. Rev. 32:38–55.

Qian W, Han Z-J, He C. 2008. Two-component signal transduction systems of Xanthomonas spp.: a lesson from genomics. Mol. Plant Microbe Interact. 21:151–161.

Rand JM, Pisithkul T, Clark RL, Thiede JM, Mehrer CR, Agnew DE, Campbell CE, Markley AL, Price MN, Ray J, et al. 2017. A metabolic pathway for catabolizing levulinic acid in bacteria. Nat. Microbiol. 2:1624–1634.

Ronneau S, Hallez R. 2019. Make and break the alarmone: regulation of (p)ppGpp synthetase/hydrolase enzymes in bacteria. FEMS Microbiol. Rev. 43:389–400.

Schmidl SR, Ekness F, Sofjan K, Daeffler KN-M, Brink KR, Landry BP, Gerhardt KP, Dyulgyarov N, Sheth RU, Tabor JJ. 2019. Rewiring bacterial two-component systems by modular DNA-binding domain swapping. Nat. Chem. Biol. 15:690–698.

Sharma P, Bano A, Singh SP, Sharma S, Xia C, Nadda AK, Lam SS, Tong YW. 2022. Engineered microbes as effective tools for the remediation of polyaromatic aromatic hydrocarbons and heavy metals. Chemosphere 306:135538.

Silva GGZ, Cuevas DA, Dutilh BE, Edwards RA. 2014. FOCUS: an alignment-free model to identify organisms in metagenomes using non-negative least squares. PeerJ 2:e425.

Simon C. 2022. An evolving view of phylogenetic support. Syst. Biol. 71:921–928.

Siryaporn A, Goulian M. 2008. Cross-talk suppression between the CpxA-CpxR and EnvZ-OmpR two-component systems in E. coli. Mol. Microbiol. 70:494–506.

Siryaporn A, Perchuk BS, Laub MT, Goulian M. 2010. Evolving a robust signal transduction pathway from weak cross-talk. Mol. Syst. Biol. 6:452.

Smith MD, Wertheim JO, Weaver S, Murrell B, Scheffler K, Kosakovsky Pond SL. 2015. Less is more: an adaptive branch-site random effects model for efficient detection of episodic diversifying selection. Mol. Biol. Evol. 32:1342–1353.

Stephenson K, Hoch JA. 2002. Evolution of signalling in the sporulation phosphorelay. Mol. Microbiol. 46:297–304.

Stock AM, Robinson VL, Goudreau PN. 2000. Two-component signal transduction. Annu. Rev. Biochem. 69:183–215.

Studier FW. 2005. Protein production by auto-induction in high density shaking cultures. Protein Expr. Purif. 41:207–234.

Thompson MG, Blake-Hedges JM, Cruz-Morales P, Barajas JF, Curran SC, Eiben CB, Harris NC, Benites VT, Gin JW, Sharpless WA, et al. 2019. Massively parallel fitness profiling reveals multiple novel enzymes in Pseudomonas putida lysine metabolism. MBio 10.

Thompson MG, Incha MR, Pearson AN, Schmidt M, Sharpless WA, Eiben CB, Cruz-Morales P, Blake-Hedges JM, Liu Y, Adams CA, et al. 2020. Fatty acid and alcohol metabolism in Pseudomonas putida: functional analysis using random barcode transposon sequencing. Appl. Environ. Microbiol. 86.

Tiwari S, Jamal SB, Hassan SS, Carvalho PVSD, Almeida S, Barh D, Ghosh P, Silva A, Castro TLP, Azevedo V. 2017. Two-Component Signal Transduction Systems of Pathogenic Bacteria As Targets for Antimicrobial Therapy: An Overview. Front. Microbiol. 8:1878.

Vandewege MW, Mangum SF, Gabaldón T, Castoe TA, Ray DA, Hoffmann FG. 2016. Contrasting patterns of evolutionary diversification in the olfactory repertoires of reptile and bird genomes. Genome Biol. Evol. 8:470–480.

Varadi M, Anyango S, Deshpande M, Nair S, Natassia C, Yordanova G, Yuan D, Stroe O, Wood G, Laydon A, et al. 2022. AlphaFold Protein Structure Database: massively expanding the structural coverage of protein-sequence space with high-accuracy models. Nucleic Acids Res. 50:D439–D444.

Varughese KI, Madhusudan, Zhou XZ, Whiteley JM, Hoch JA. 1998. Formation of a novel four-helix bundle and molecular recognition sites by dimerization of a response regulator phosphotransferase. Mol. Cell 2:485–493.

Volke DC, Turlin J, Mol V, Nikel PI. 2020. Physical decoupling of XylS/Pm regulatory elements and conditional proteolysis enable precise control of gene expression in Pseudomonas putida. Microb. Biotechnol. 13:222–232.

Wang BX, Cady KC, Oyarce GC, Ribbeck K, Laub MT. 2021. Two-Component Signaling Systems Regulate Diverse Virulence-Associated Traits in Pseudomonas aeruginosa. Appl. Environ. Microbiol. 87.

Weaver S, Shank SD, Spielman SJ, Li M, Muse SV, Kosakovsky Pond SL. 2018. Datamonkey 2.0: A modern web application for characterizing selective and other evolutionary processes. Mol. Biol. Evol. 35:773–777.

Wetmore KM, Price MN, Waters RJ, Lamson JS, He J, Hoover CA, Blow MJ, Bristow J, Butland G, Arkin AP, et al. 2015. Rapid quantification of mutant fitness in diverse bacteria by sequencing randomly bar-coded transposons. MBio 6:e00306–15.

Zhang Z, Wood WI. 2003. A profile hidden Markov model for signal peptides generated by HMMER. Bioinformatics 19:307–308.

