## Supplementary figures and images for "REC protein family expansion by the emergence of a new signaling pathway"

### ABSREL_model12.png

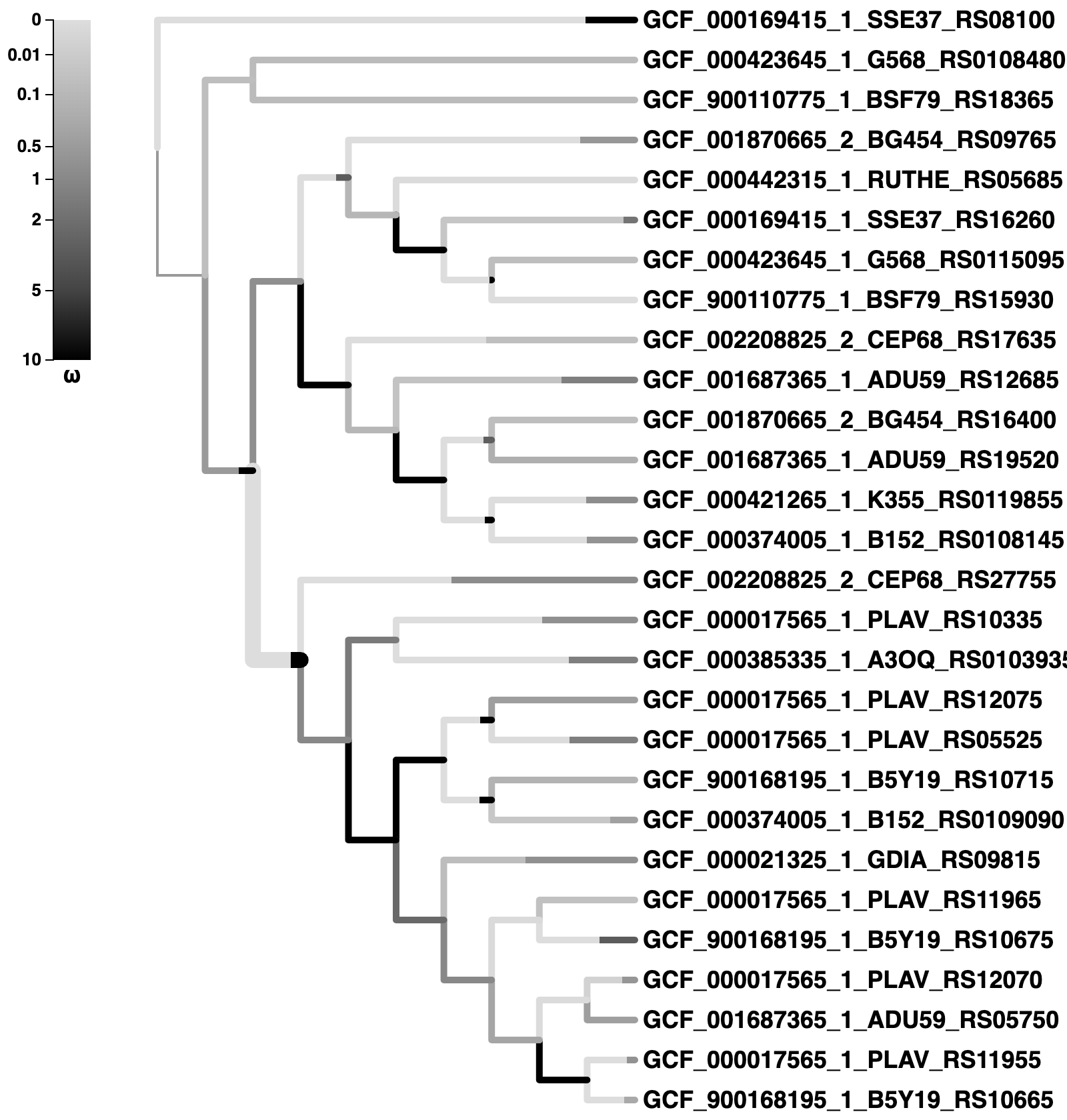

### MEME_alpha.png

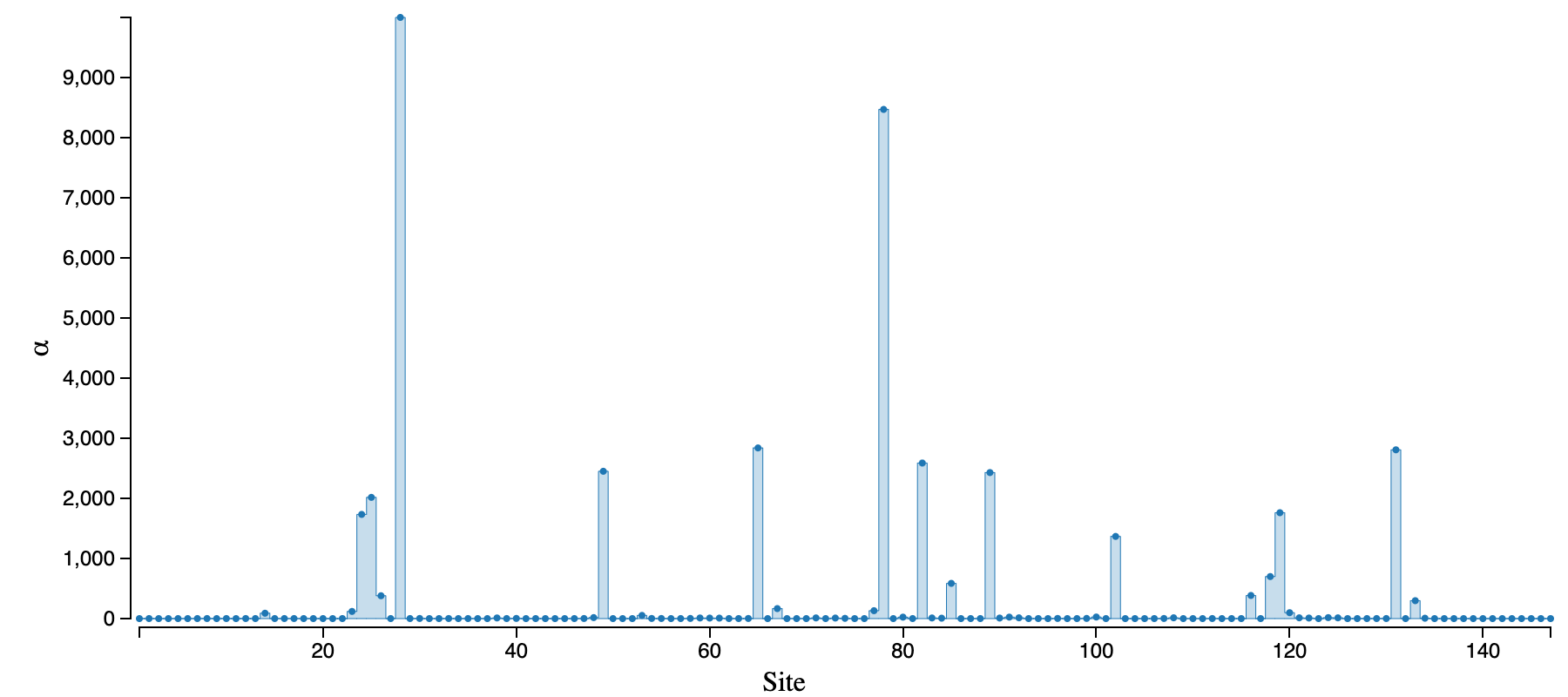

### MEME_pval.png

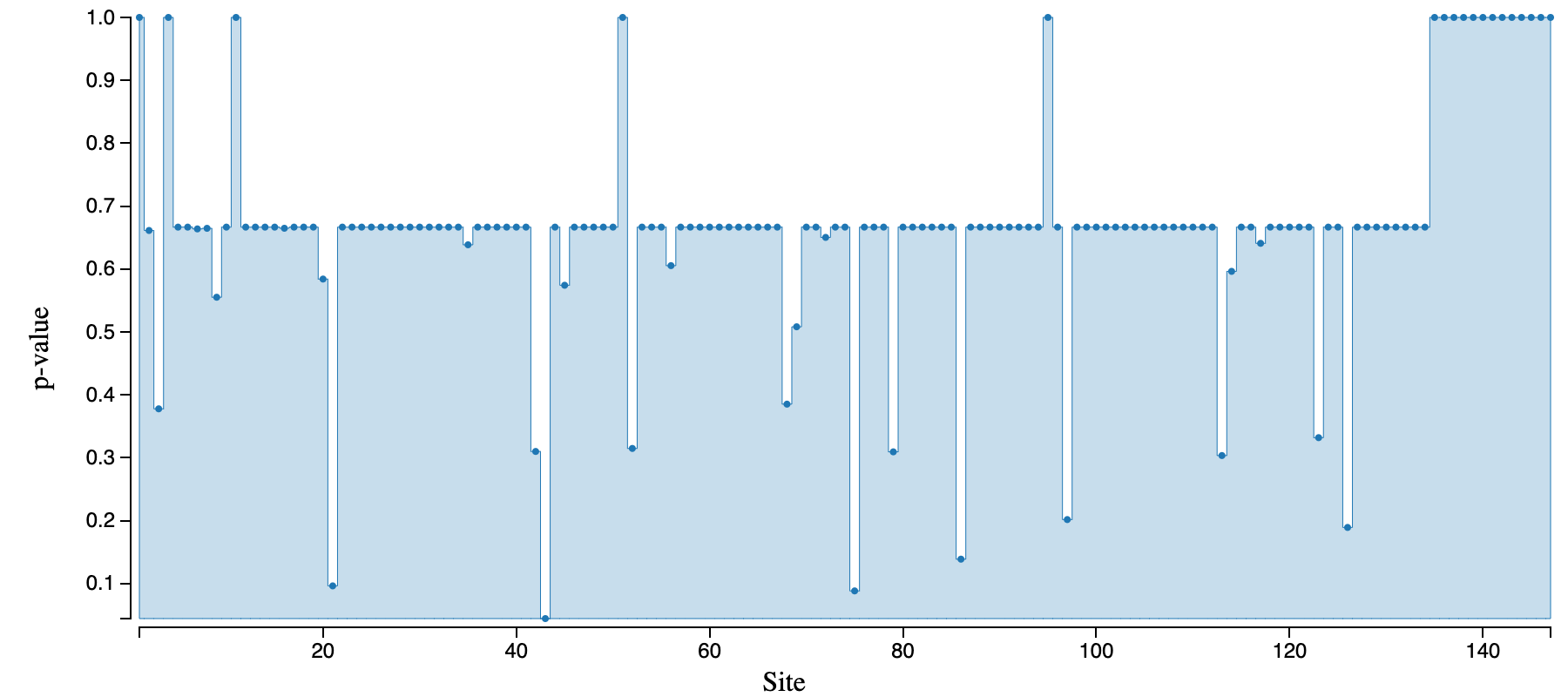

### PP1066_full_b5392_coverage.png

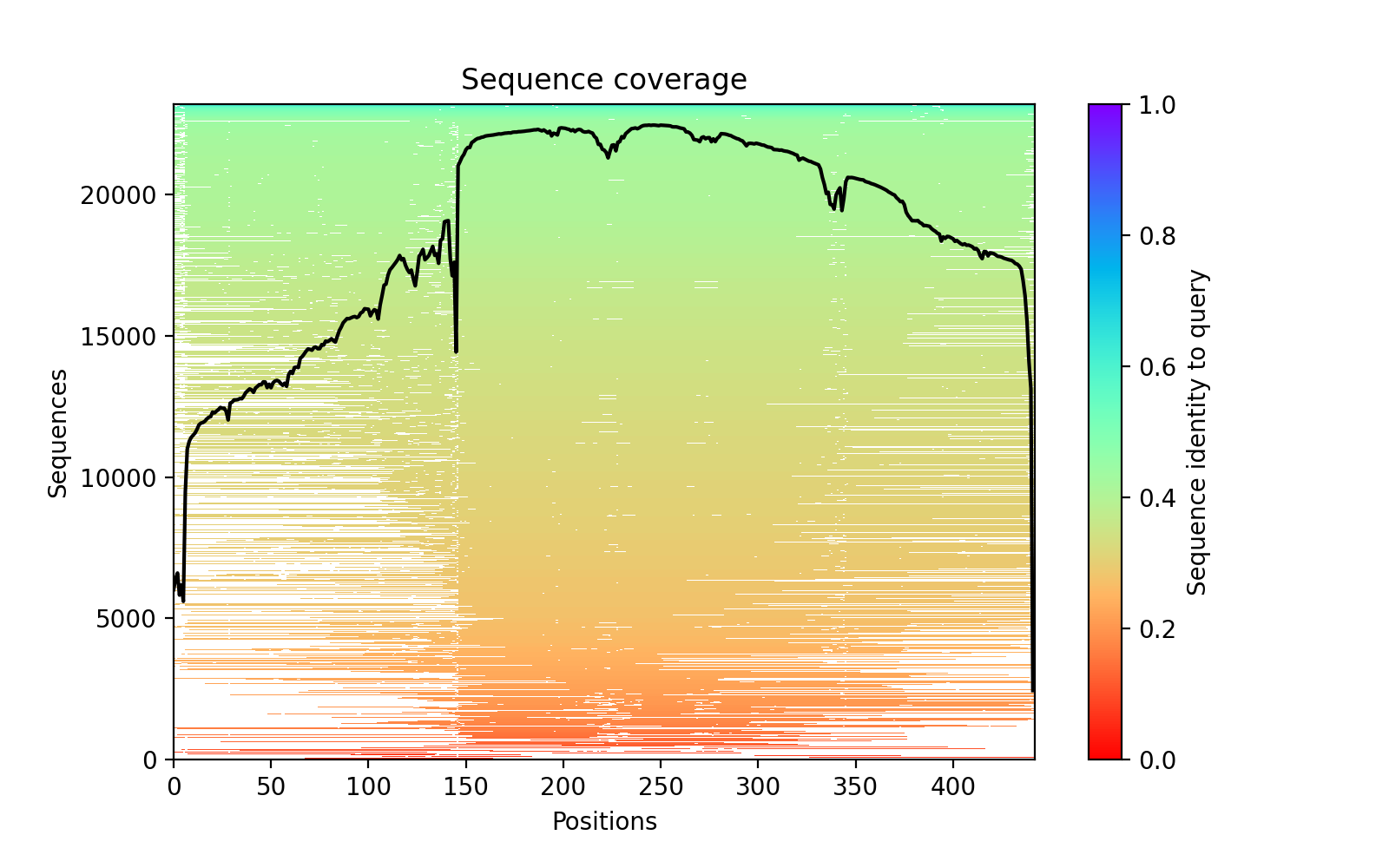

### PP1066_full_b5392_PAE.png

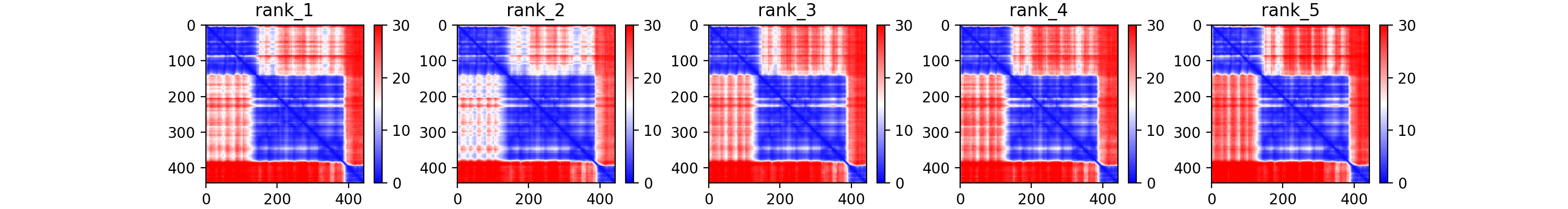

### PP1066_full_b5392_plddt.png

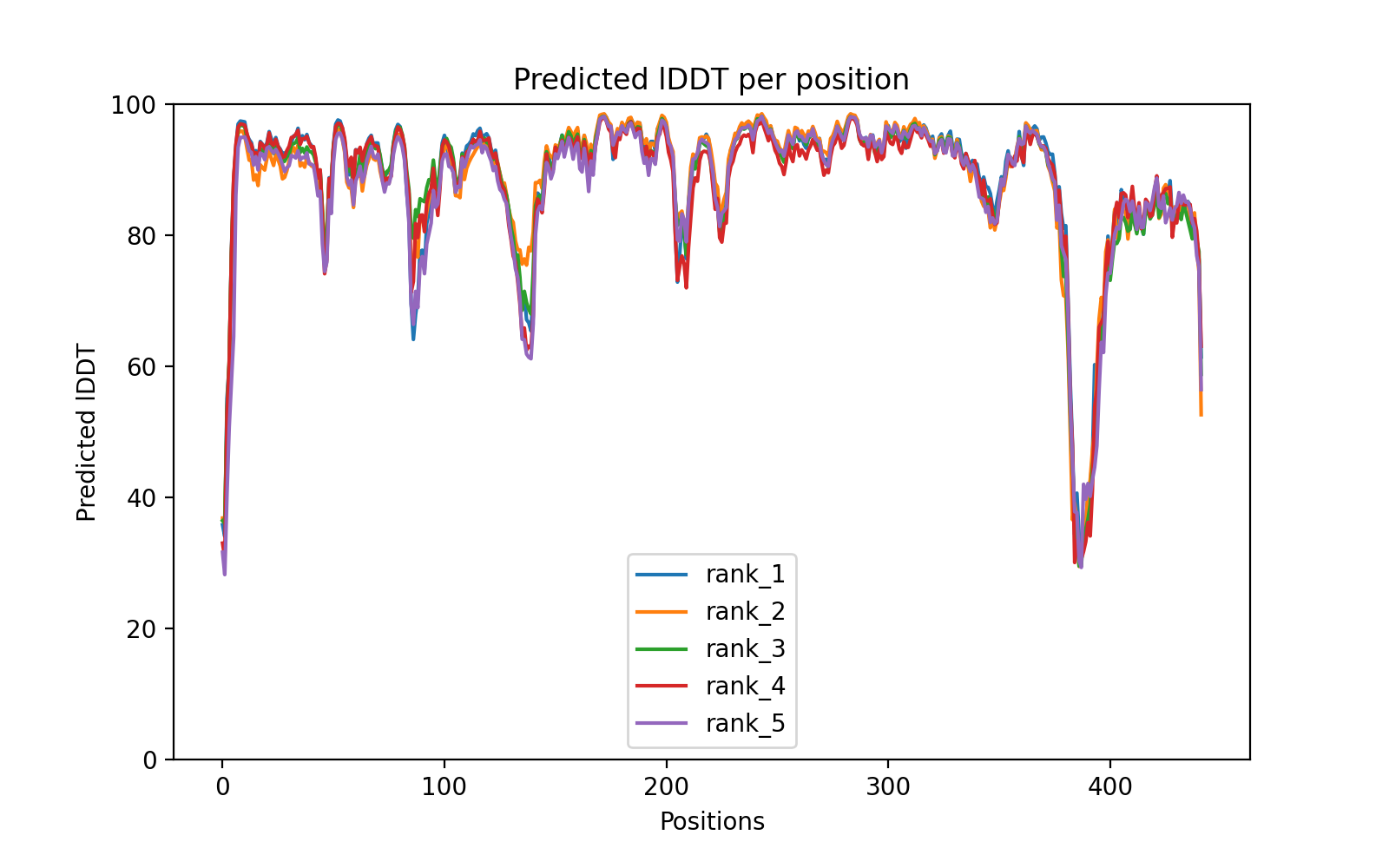

### PP1066_mut_0d0be_coverage.png

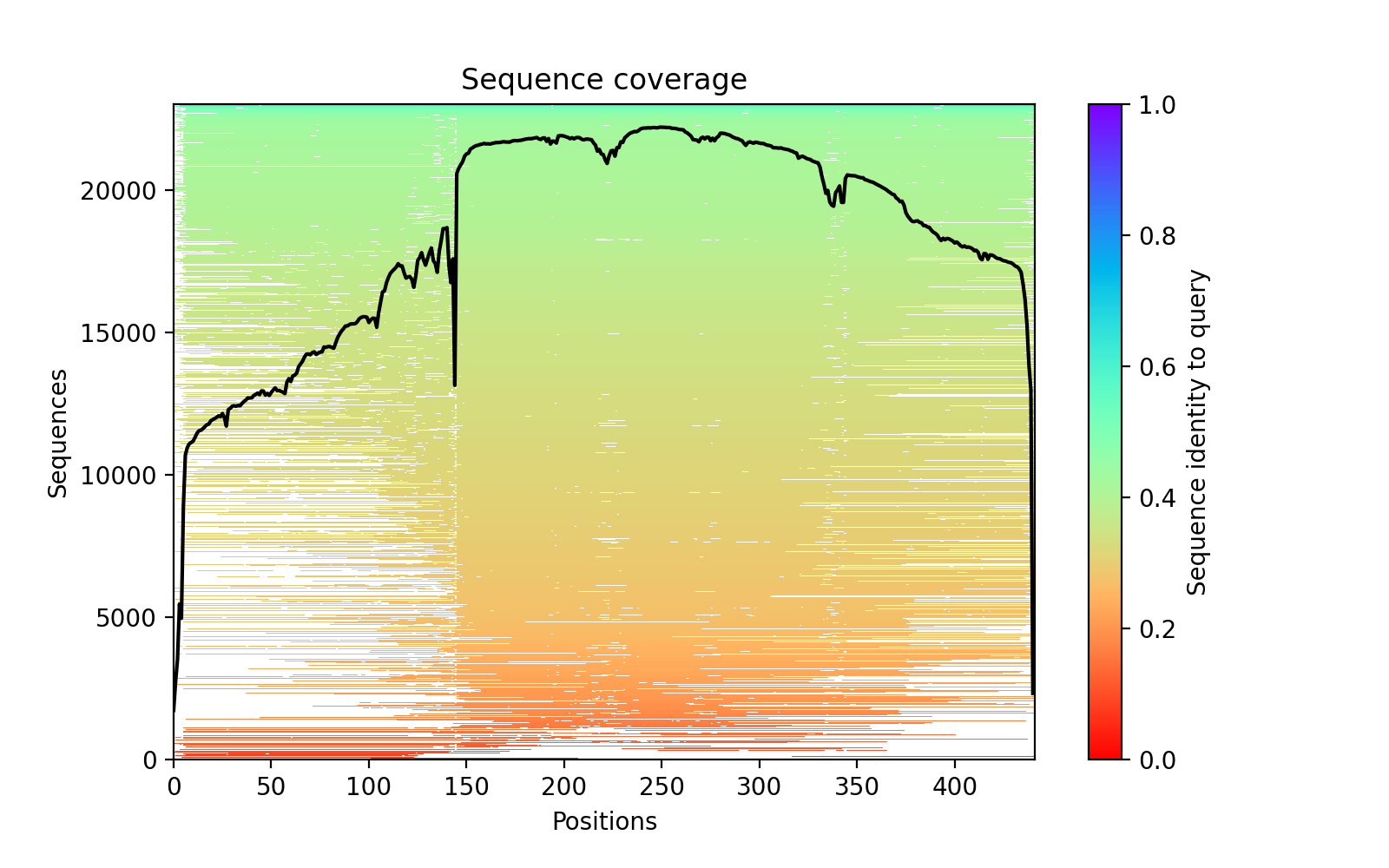

### PP1066_mut_0d0be_PAE.png

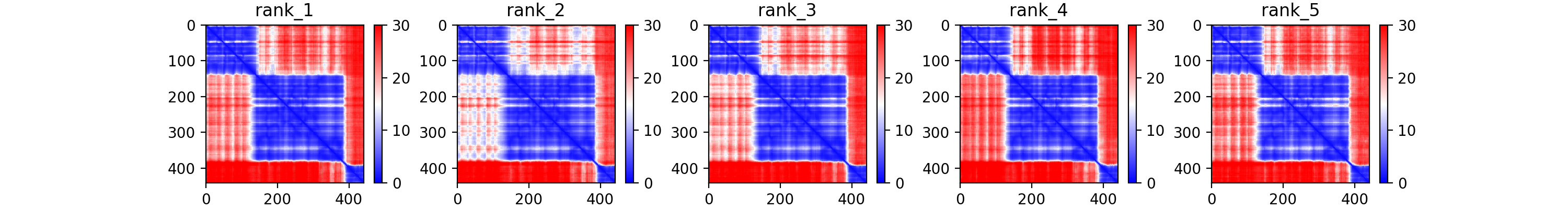

### PP1066_mut_0d0be_plddt.png

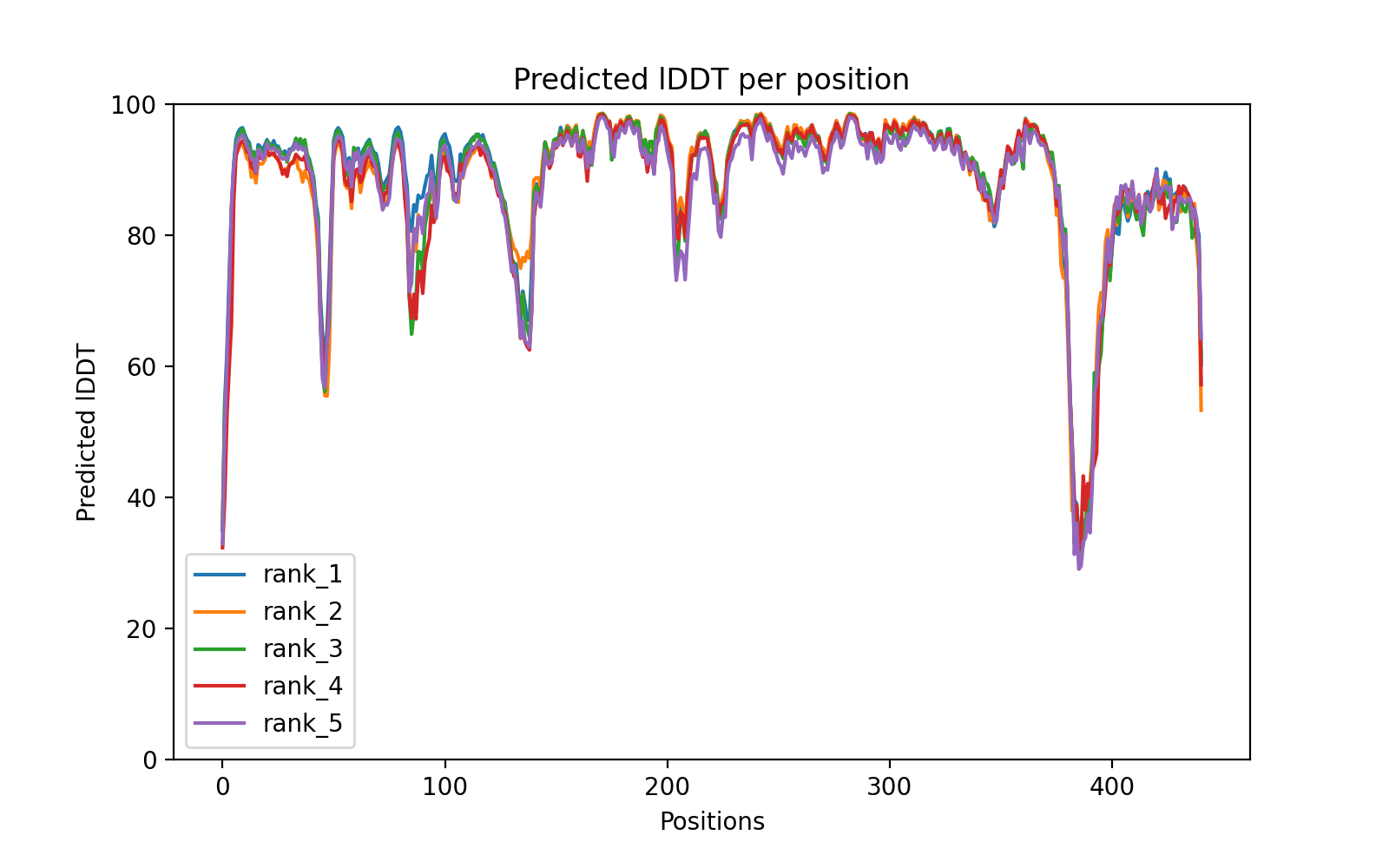

### PP3551_mut_d0b08_coverage.png

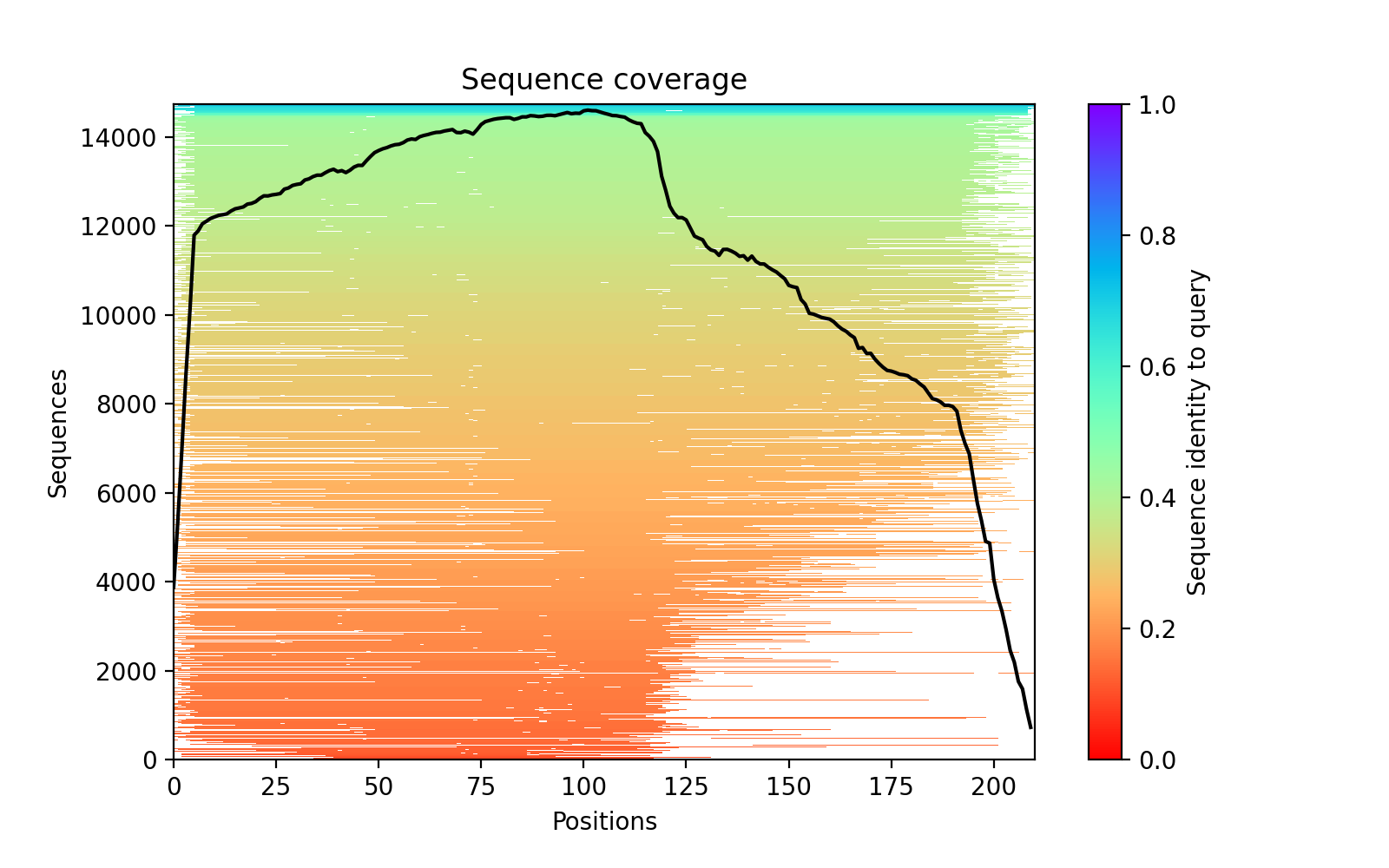

### PP3551_mut_d0b08_PAE.png

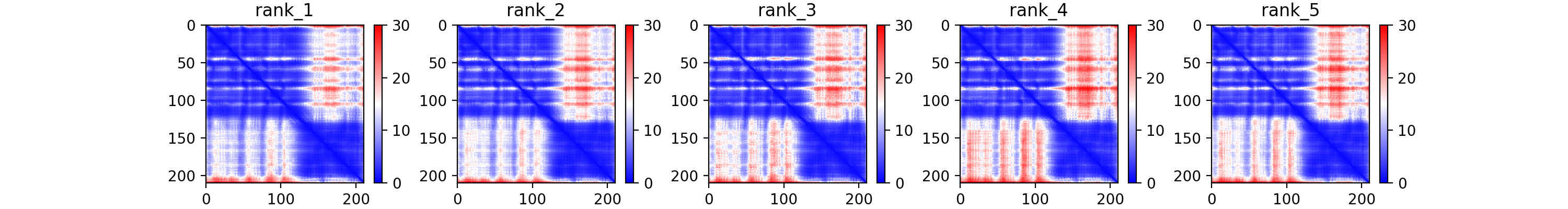

### PP3551_mut_d0b08_plddt.png

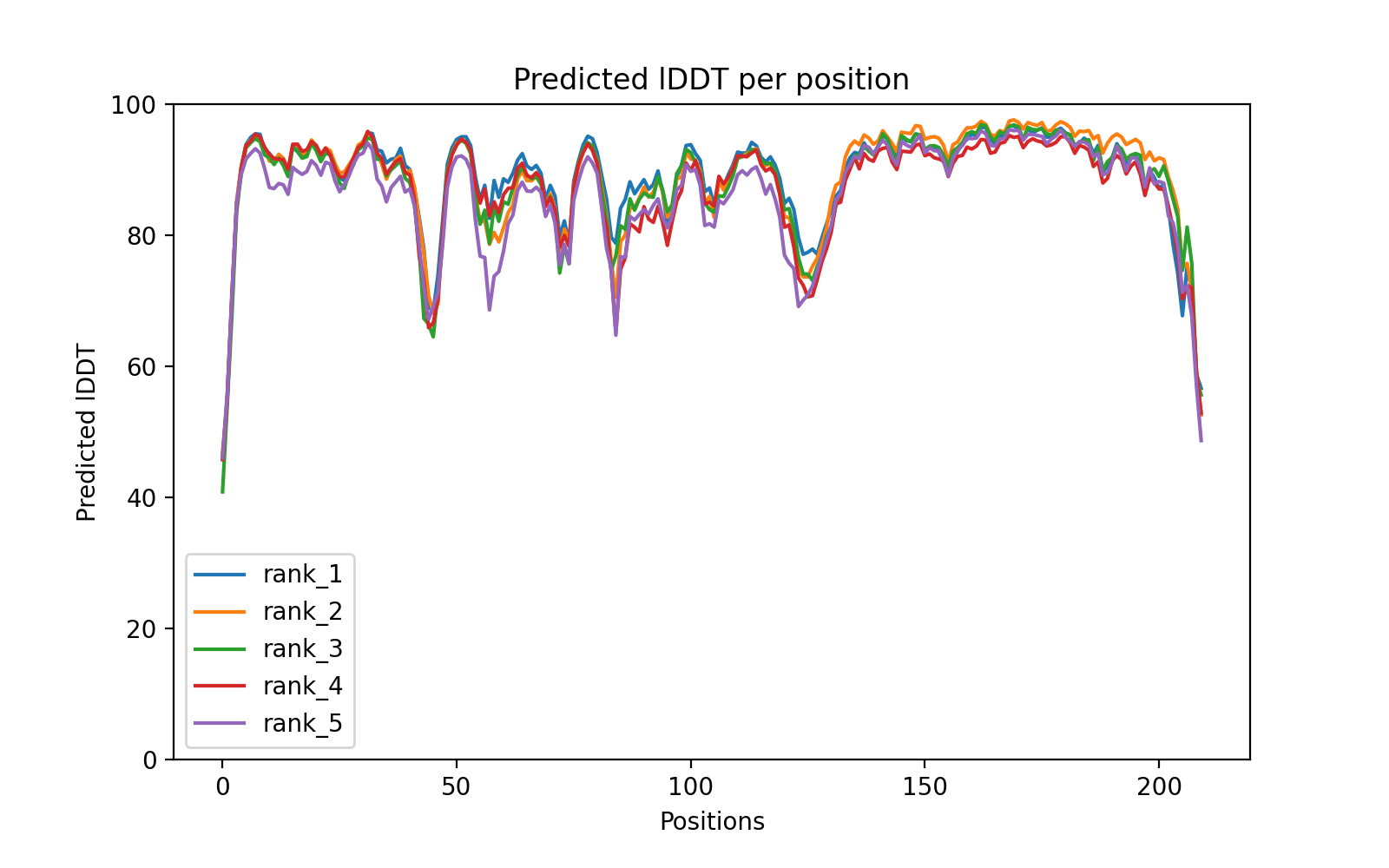
