## Supplemental Data 1 for "REC protein family expansion by the emergence of a new signaling pathway": MEME_log.txt

Analysis Description
--------------------
MEME (Mixed Effects Model of Evolution) estimates a site-wise synonymous
(&alpha;) and a two-category mixture of non-synonymous (&beta;-, with
proportion p-, and &beta;+ with proportion [1-p-]) rates, and uses a
likelihood ratio test to determine if &beta;+ > &alpha; at a site. The
estimates aggregate information over a proportion of branches at a site,
so the signal is derived from episodic diversification, which is a
combination of strength of selection [effect size] and the proportion of
the tree affected. A subset of branches can be selected for testing as
well, in which case an additional (nuisance) parameter will be inferred
-- the non-synonymous rate on branches NOT selected for testing.
Multiple partitions within a NEXUS file are also supported for
recombination - aware analysis. 

- __Requirements__: in-frame codon alignment and a phylogenetic tree

- __Citation__: Detecting Individual Sites Subject to Episodic Diversifying Selection.
_PLoS Genet_ 8(7): e1002764.

- __Written by__: Sergei L. Kosakovsky Pond, Steven Weaver

- __Contact Information__:

- __Analysis Version__: 2.1.2


>code –> Universal

>Loaded a multiple sequence alignment with **28** sequences, **147** codons, and **1** partitions from `/home/datamonkey/datamonkey-js-server/production/app/meme/output/631cd621fcc5751437bd1459`

>branches –> All

>Select the p-value threshold to use when testing for selection (permissible range = [0,1], default value = 0.1): 
>pvalue –> 0.1

>[Advanced setting, will result in MUCH SLOWER run time] Perform parametric bootstrap resampling to derive site-level null LRT distributions up to this many replicates per site. Recommended use for small to medium (<30 sequences) datasets (permissible range = [0,1000], default value = 50, integer): 
>resample –> 0


### Branches to include in the MEME analysis
Selected 53 branches to include in the MEME analysis: `GCF_000021325_1_GDIA_RS09815, GCF_000017565_1_PLAV_RS12070, GCF_001687365_1_ADU59_RS05750, Node5, GCF_000017565_1_PLAV_RS11955, GCF_900168195_1_B5Y19_RS10665, Node8, Node4, GCF_000017565_1_PLAV_RS11965, GCF_900168195_1_B5Y19_RS10675, Node11, Node3, Node1, GCF_000423645_1_G568_RS0108480, GCF_900110775_1_BSF79_RS18365, Node18, GCF_000169415_1_SSE37_RS08100, Node17, GCF_000423645_1_G568_RS0115095, GCF_900110775_1_BSF79_RS15930, Node26, GCF_000169415_1_SSE37_RS16260, Node25, GCF_000442315_1_RUTHE_RS05685, Node24, GCF_001870665_2_BG454_RS09765, Node23, GCF_002208825_2_CEP68_RS17635, GCF_001687365_1_ADU59_RS12685, GCF_001870665_2_BG454_RS16400, GCF_001687365_1_ADU59_RS19520, Node37, GCF_000421265_1_K355_RS0119855, GCF_000374005_1_B152_RS0108145, Node40, Node36, Node34, Node32, Node22, Node16, GCF_002208825_2_CEP68_RS27755, Node15, GCF_000017565_1_PLAV_RS10335, GCF_000385335_1_A3OQ_RS0103935, Node44, Node14, GCF_000017565_1_PLAV_RS12075, GCF_000017565_1_PLAV_RS05525, Node48, GCF_900168195_1_B5Y19_RS10715, GCF_000374005_1_B152_RS0109090, Node51, Node47`


### Obtaining branch lengths and nucleotide substitution biases under the nucleotide GTR model

>kill-zero-lengths –> Yes
* Log(L) = -8934.85, AIC-c = 17992.31 (61 estimated parameters)
* 1 partition. Total tree length by partition (subs/site)  9.738

### Obtaining the global omega estimate based on relative GTR branch lengths and nucleotide substitution biases
* Log(L) = -8480.19, AIC-c = 17098.70 (68 estimated parameters)
* 1 partition. Total tree length by partition (subs/site) 16.349
* non-synonymous/synonymous rate ratio for *test* =   0.1498

### Improving branch lengths, nucleotide substitution biases, and global dN/dS ratios under a full codon model

>full-model –> Yes
* Log(L) = -8390.81
* non-synonymous/synonymous rate ratio for *test* =   0.0246

### For partition 1 these sites are significant at p <=0.1

|   Codon    | Partition  |   alpha    |   beta+    |     p+     |    LRT     |Episodic selection detected?| # branches |Most common codon substitutions at this site|
|:----------:|:----------:|:----------:|:----------:|:----------:|:----------:|:--------------------------:|:----------:|:------------------------------------------:|
|     21     |     1      |    0.000   |    0.096   |    0.456   |    3.196   |      Yes, p =  0.0963      |     2      |[2]AAG>ATG|[1]AAG>ACG,AAG>AGC,AAG>ATC,AAG...|
|     43     |     1      |    0.039   |    0.457   |    0.146   |    4.686   |      Yes, p =  0.0443      |     1      |[2]CTG>CTC,CTG>TTG,TGG>ACG|[1]ATG>TGG,CTG...|
|     75     |     1      |    0.015   |    1.123   |    0.149   |    3.360   |      Yes, p =  0.0884      |     0      |[2]ACC>AGC,ACC>TCC|[1]ACC>GCG,ACC>GTC,ACC...|
