## Supplementary material for "REC protein family expansion by the emergence of a new signaling pathway": Supplmental Materials

### Supplemental Materials for Garber *et al.*

#### Document Contents:

**Supplemental Figures 1-10**

**Supplemental Tables 1-4**

**Supplemental Methods**

**Supplemental Data 1.**
Results log from DataMonkey version of HyPhy's absREL and MEME. Contents include original fasta, log.txt, result.json, results tree, results table (if applicable), and results summary figure.

**Supplemental Data 2.**
Results log and output from ColabFold for PP_1066, PP_1066* (PP_1066_mut) and PP3551* (PP_3551_mut).

**Supplemental Data 3.**
Results output from motif calling with orthologous RRs. Each tab contains data for each ortholog.

**Supplemental Data 4.**
Raw output from DAP-seq peak calling

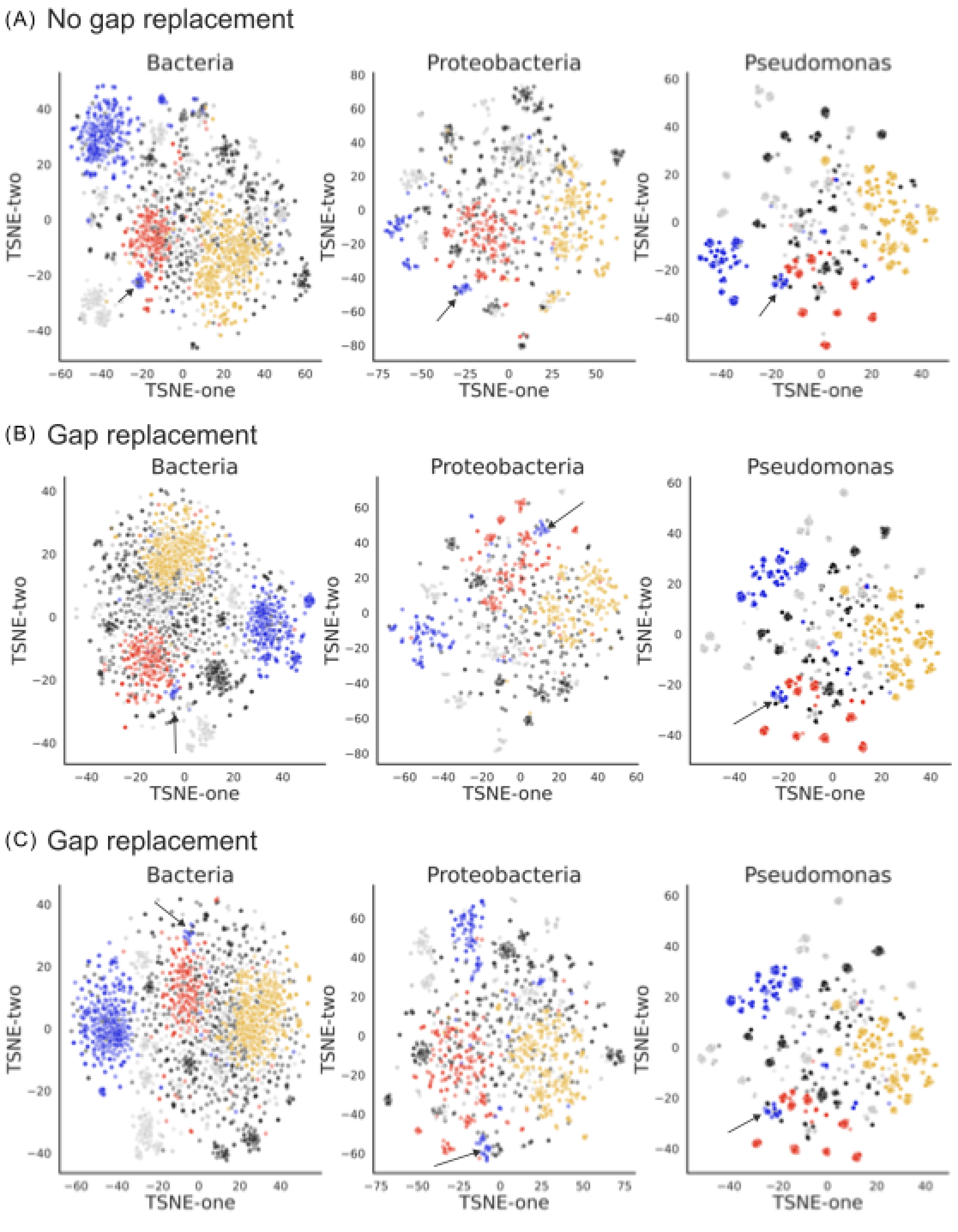


Supplemental Figure 1

***Supplemental Figure 1: Dimensionally reduced REC domain sequence alignments with and without gap replacement show similar data structures.***

Output from the machine learning algorithm assesses the similarity between members of the REC protein family in Bacteria (leftmost panel), Proteobacteria (middle panel), and Pseudomonas (right panel) species, showing the relationship between REC domain sequence alignments with the same data as Figure 1D but without gap replacement (A) and with independently sampled data with gap replacement

(B, C) and explaining 95% of the variation between the sequences. The REC domains were sampled from species found within the taxonomic rank (kingdom - Bacteria, phylum-Proteobacteria, or genera-Pseudomonas) labeled at the top of each plot. To control for overrepresentation of subclades due to more representative species with full genome sequences in any given subclade, species were sampled evenly from the taxonomic rank below the one represented in the plot title (e.g. The taxonomic rank below kingdom is phylum; the maximum of species from each bacterial phylum were randomly sampled to generate the bacteria dataset (see Table S1)). After species sampling, we identified all proteins with REC domains in each of the sampled species' genomes and aligned their sequences. To read the plot, each point represents the sequence of a REC domain from the sampled genomes, points that are close together share sequence identity with each other. Each REC domain is annotated by the identity of its effector domain Trans_reg_C (yellow) , GerE (blue), AAA+ (red), CheY (gray), other (gray outline), and no domains (black). Note that the location of REC domains is not fixed in the plot between independent runs of the algorithm, however, the relative distribution between the points is visually consistent between runs. The within-gene-recombination that changed parent REC domain fused to AAA+ domain to REC domain fused to GerE domain is apparent by the presence of two distinct blue clusters: the GerE cluster (blue) found close to the AAA+ cluster (red) resulted from within-gene-recombination of effector domains and is indicated by a black arrow.

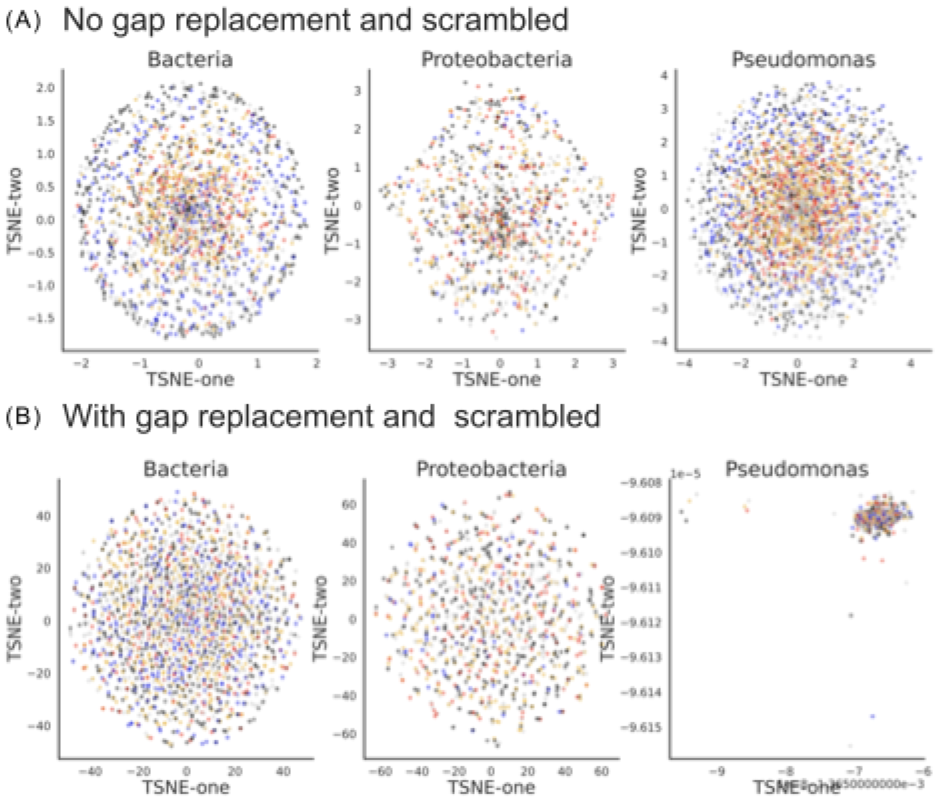


Supplemental Figure 2

***Supplemental Figure 2: Dimensionally reduced of scrambled REC domain sequence alignments with and without gap replacement reveals bias from the alignment strategy.***

Output from the dimensional reduction algorithm assesses the similarity between members of the REC protein family in Bacteria (leftmost panel), Proteobacteria (middle panel), and Pseudomonas (right panel) species, showing the relationship between scrambled REC domain sequence alignments with the same data as Figure 1D but without gap replacement (A) with gap replacement (B) and explaining 95% of the variation between the sequences. The REC domains were sampled from species found within the taxonomic rank (kingdom - Bacteria, phylum-Proteobacteria, or genera-Pseudomonas) labeled at the top of each plot. To control for overrepresentation of subclades due to more representative species with full genome sequences in any given subclade, species were sampled evenly from the taxonomic rank below the one represented in the plot title (e.g. The taxonomic rank below kingdom is phylum; the maximum of species from each bacterial phylum were randomly sampled to generate the bacteria dataset (see Table S1)). After species sampling, we identified all proteins with REC domains in each of the sampled species' genomes and aligned their sequences. To read the plot, each point represents the sequence of a REC domain from the sampled genomes, points that are close together share sequence identity with each other. Each REC domain is annotated by the identity of its effector domain Trans_reg_C (yellow) , GerE (blue), AAA+ (red), CheY (gray), other (gray outline), and no domains (black). Note that the location of REC domains is not fixed in the plot between independent runs of the algorithm, however, the relative distribution between the points is visually consistent between runs. The within-gene-recombination that changed parent REC domain fused to AAA+ domain to REC domain fused to GerE domain is apparent by the presence of two distinct blue clusters: the GerE cluster (blue) found close to the AAA+ cluster (red) resulted from within-gene-recombination of effector domains and is indicated by a black arrow.

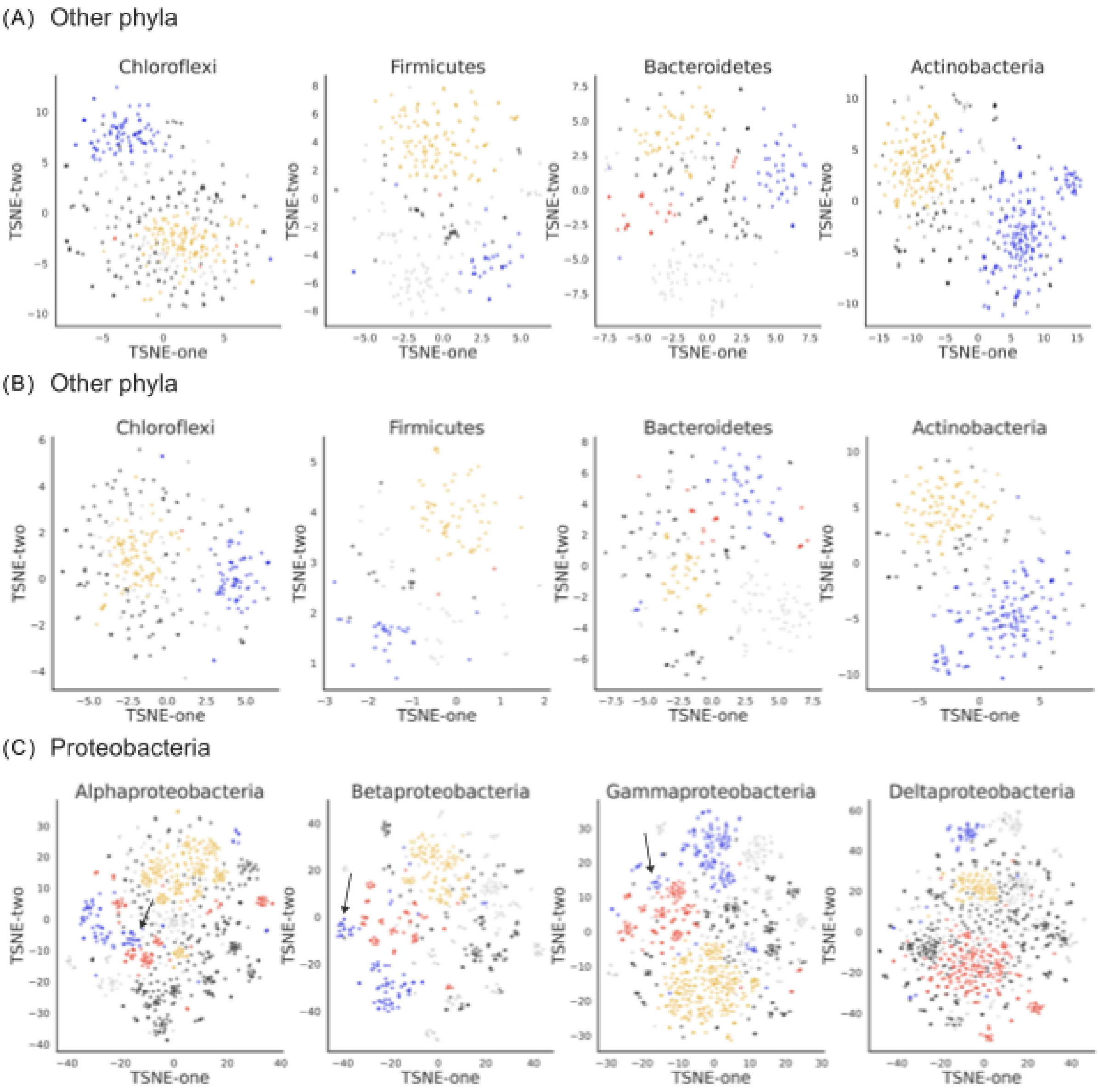


Supplemental Figure 3

***Supplemental Figure 3: Within-gene-recombination that changed parent REC domain fused to AAA+ domain to REC domain fused to GerE domain occurred during the Proteobacteria lineage, specifically the Alphaproteobacteria lineage.***

(A, B) Output from the machine learning algorithm assesses the similarity between members of the REC protein family from two independently sampled datasets (shown respectively in A and B) in (from left to right) Chloroflexi, Firmicutes, Bacteroidetes, Actinobacteria species, showing the relationship between REC domain sequence alignments with gap replacement and explaining 95% of the variation between the sequences. The REC domains were sampled from species found within the taxonomic rank (phylum) labeled at the top of each plot. To control for overrepresentation of subclades due to more representative species with full genome sequences in any given subclade, species were sampled evenly from the taxonomic rank below the one represented in the plot title (e.g. the taxonomic rank below phylum is class; the maximum species from each of the phylum's classes were randomly sampled to generate the phylum dataset). After species sampling, we identified all proteins with REC domains in each of the sampled species' genomes and aligned their sequences. To read the plot, each point represents the sequence of a REC domain from the sampled genomes, points that are close together share sequence identity with each other. Each REC domain is annotated by the identity of its effector domain Trans_reg_C (yellow) , GerE (blue), AAA+ (red), CheY (gray), other (gray outline), and no domains (black). Note that the location of REC domains is not fixed in the plot between independent runs of the algorithm, however, the relative distribution between the points is visually consistent between runs. The within-gene-recombination that changed parent REC domain fused to AAA+ domain to REC domain fused to GerE domain is apparent by the presence of two distinct blue clusters: the GerE cluster (blue) found close to the AAA+ cluster (red) resulted from within-gene-recombination of effector domains and is indicated by a black arrow.

(B) Output from the machine learning algorithm assesses the similarity between members of the REC protein family from an independently sampled dataset compared to Figure 2 in (from left to right) Alphaproteobacteria, Betaproteobacteria, Gammaproteobacteria, Deltaproteobacteria species, showing the relationship between REC domain sequence alignments with gap replacement and explaining 95% of the variation between the sequences. The REC domains were sampled from species found within the taxonomic rank (class) labeled at the top of each plot. To control for overrepresentation of subclades due to more representative species with full genome sequences in any given subclade, species were sampled evenly from the taxonomic rank below the one represented in the plot title (e.g. the taxonomic rank below class is order; the maximum of species from each bacterial phylum were randomly sampled to generate the bacteria dataset (see Table S1)). After species sampling, we identified all proteins with REC domains in each of the sampled species' genomes and aligned their sequences. To read the plot, each point represents the sequence of a REC domain from the sampled genomes, points that are close together share sequence identity with each other. Each REC domain is annotated by the identity of its effector domain Trans_reg_C (yellow) , GerE (blue), AAA+ (red), CheY (gray), other (gray outline), and no domains (black). Note that the location of REC domains is not fixed in the plot between independent runs of the algorithm, however, the relative distribution between the points is visually consistent between runs. The within-gene-recombination that changed parent REC domain fused to AAA+ domain to REC domain fused to GerE domain is apparent by the presence of two distinct blue clusters: the GerE cluster (blue) found close to the AAA+ cluster (red) resulted from within-gene-recombination of effector domains and is indicated by a black arrow.

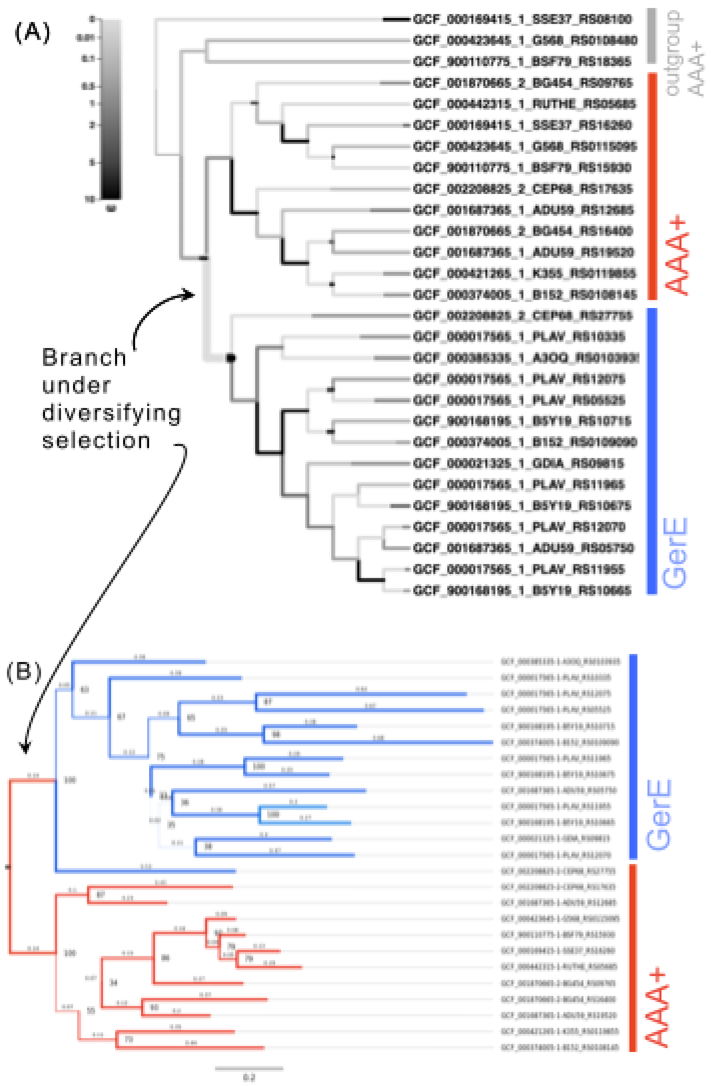


Supplemental Figure 4

***Supplemental Figure 4:***

(A) The REC domains within the black box in Figure 3A were used to generate a REC domain DNA sequence alignment with MAFFT to test for episodic diversification on the branches separating the parent REC domains with AAA+ effectors (red) and recombined REC domains with GerE (or no domain) effectors (blue). Episodic diversification occurred after the within-gene-recombination (thick gray branch).

(B) The REC domains within the black box (Figure 3A), excluding outgroup domains, are used to construct a domain tree. The DNA sequences of the REC domains were aligned with MAFFT. We built the representative domain tree with iqtree, and annotated the branch colors red if the domain's effector identity was AAA+ or blue if the domain's effector identity was GerE (or no domain). Bootstrap supports (labeled at the nodes and shown as branch thickness) are 100 for the nodes separating the parent REC domains with AAA+ effectors and recombined REC domains with GerE (or no domain) effectors. We independently determined that the recombined REC domains with GerE (or no domain) effectors were under episodic diversifying selection. Here and Figure 3B, the parent REC domains with AAA+ effectors (red) independently duplicated and diverged after the within-gene-recombination. Sequence alignments in Figure 3C,D and Figure 4C show the parent REC domains differ at alignment residue 16 having either alanine (PP1066), tryptophan (PP1401), or threonine (PP0263) at that position. From these trees, we propose that PP1066 and its orthologs have the ancestral parent REC domain with alanine at alignment position 16.

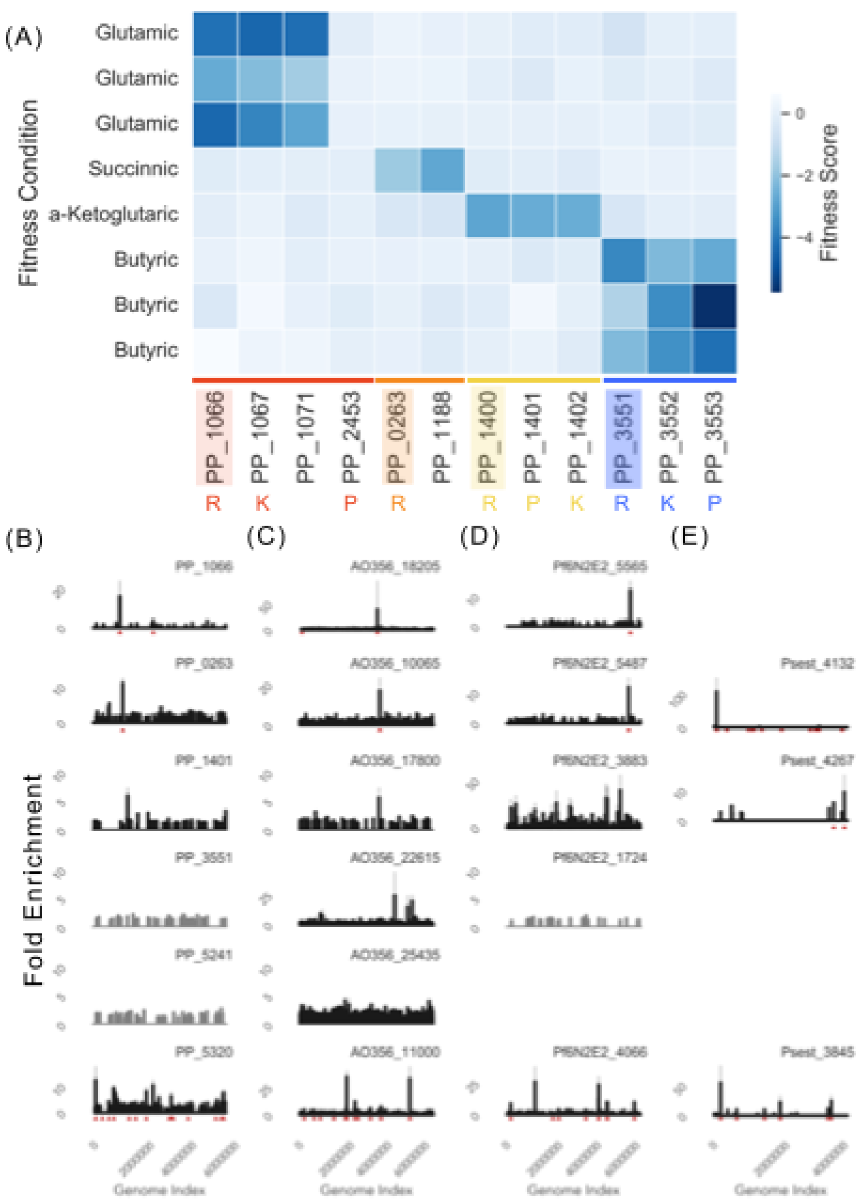


Supplemental Figure 5

***Supplemental Figure 5:***

(A) A pool of randomly barcoded transposon (RB-Tn) insertions were grown in defined media with glutamic, succinic, alpha-ketoglutaric, or butyric acids as the sole carbon source. We show the fitness score (average fold change of RB-Tn insertions within a given gene before and after competition in a pooled experiment) for genes coding for the response regulators in *P. putida* KT2440 that have the parent REC domains with AAA+ effectors (PP1066, PP0263, PP1401) and response regulator with the recombined REC domain (PP3551) when the pool was grown in the carbon-source. Although not discussed in the main text, PP1401 and PP0263 are post-duplication paralogs to PP1066. We also show fitness scores for genes related to the response regulators by synteny, co-fitness, or the presence of a DNA binding site. The function of genes are annotated below the gene ID: R = response regulator, K = histidine kinase, P = promoter (this is the promoter region used in the GFP reporter assay). (B) We applied DNA affinity purification with next generation sequencing to orthologs of the response regulators

(PP1066, PP1401, PP0263, PP3551, PP5241, and PP5320 (PhoB) in other three *Pseudomonas* species (*P. stutzeri* RCH2 (Psest), *P. fluorescens* N2E2 (Pf6N2E2), and *P. fluorescens* N2C3 (AO356). Fold enrichment of each response regulator plotted as a function of the respective genome location (Genome Index). The plot is organized by grouping together orthologous RRs along the x-axis, and by grouping RRs with paralogous REC domains along the y-axis. Red triangles below each individual plot indicate the genomic location of a binding motif. Samples that failed to enrich DNA are shown in gray. The bottom panels of the plot show data for the well-studied, positive control, PhoB. Genes with higher than background fold-enrichment are summarized in Table S2.

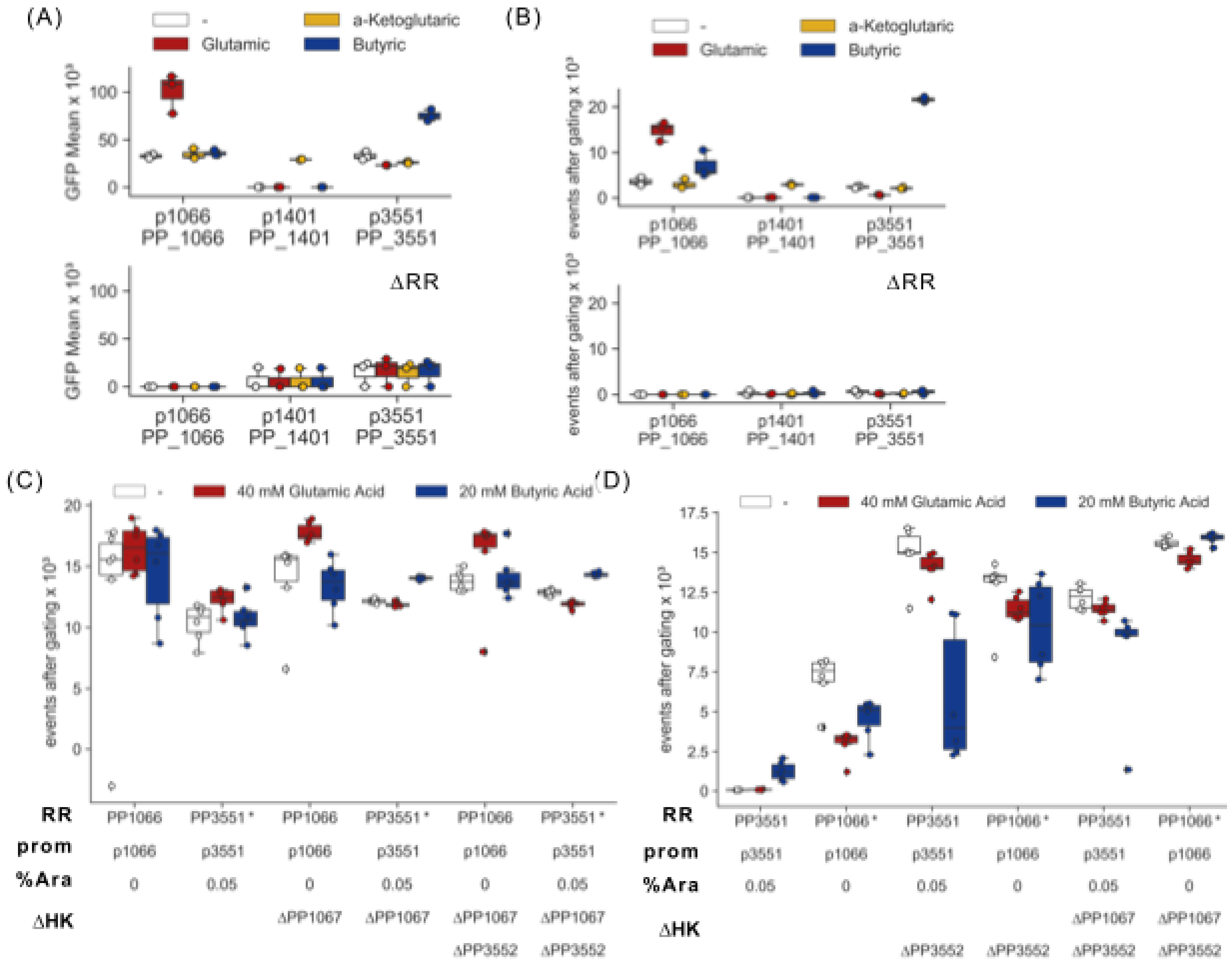


Supplemental Figure 6

***Supplemental Figure 6:***
(A) We confirmed the model for regulation by carboxylic acids with GFP reporter screens. *P. putida* KT2440 transformed with a GFP reporter plasmid for regulation by PP1066, PP1401 or PP3551 were grown in defined glucose media with or without (white) glutamic acid (red), alpha-ketoglutaric (yellow) or butyric acid (blue). Although not discussed in the main text, PP1401 is a post-duplication paralog to PP1066.Top panel shows the reporter plasmid for PP1066 is upregulated when grown with glutamic acid, PP1401 is upregulated when grown with alpha-ketoglutaric acid and the reporter plasmid for PP3551 is upregulated when grown with butyric acid. Bottom panel shows reporters for PP1066, PP1401, and PP3551 are respectively not activated if the regulators, PP1066, PP1401, or PP3551 are deleted from the genome. Fluorescence was measured by flow cytometry and measurements are reported as GFP Mean x $10^3$after gating. (B,C,D) Number of events after gating for Supplementary Figure 6A (B), Figure 4I (C) and Figure 4J (D). Results show the number of events after gating included in the calculation for GFP mean. If the number of events was < 150, we excluded the data and set the GFP mean to 0. Center line, median; box limits, upper and lower quartiles; whiskers, 1.5x interquartile range; points with black diamonds, outliers; n = 3(A,B) and n=6 (C,D).

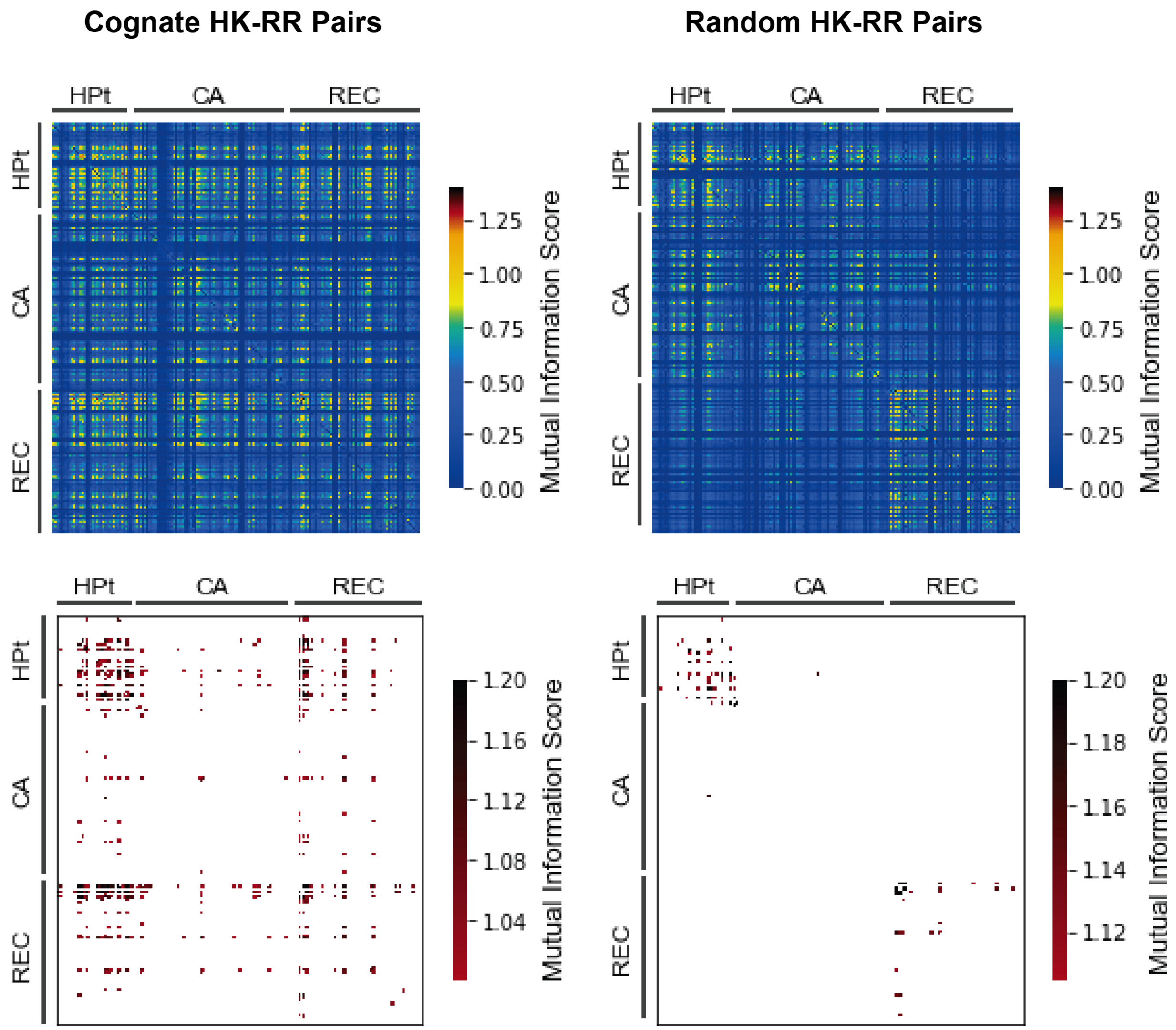


***Supplemental Figure 7:***
Covariant residues used in specificity switching assay (see Figure 4D-I) were identified by mutual information scoring of residues between REC domains and DHp-CA domains in pairs of two-component systems (left panels). Pairs of REC and DHp-CA domains were identified by syntenic relationship to each other in their respective genomes. Paired REC and DHp-CA domains show >1 mutual information scores (bottom panels) at 7 positions in the REC domain, these positions are covariant with DHp-CA domains and are interpreted as biochemically relevant for interaction and specificity of the REC domain. Randomly shuffled pairs of REC and DHp-CA domains do not show mutual information scores above the cutoff (right panels).

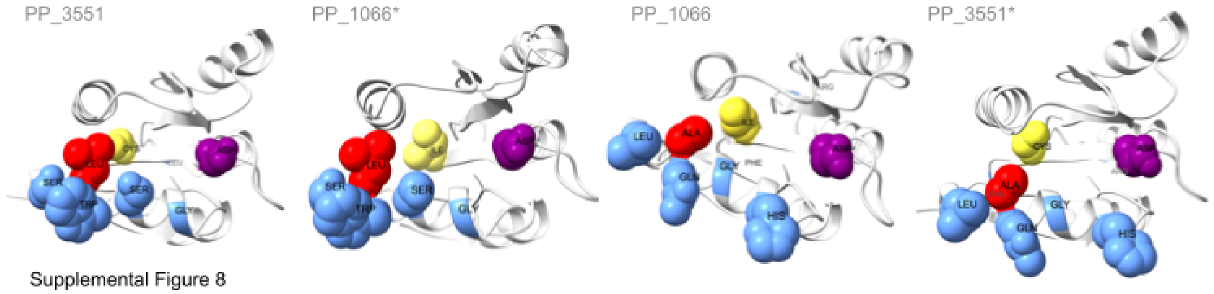


Supplemental Figure 8

***Supplemental Figure 8:***
To show the structural impact of specificity switching, we modeled the structure of PP3551, PP1066, and their respective mutant with AlphaFold. Covariant (blue), selected (yellow), covariant and selected (red), active aspartate (purple) residues are highlighted and shown in a space-filling model.

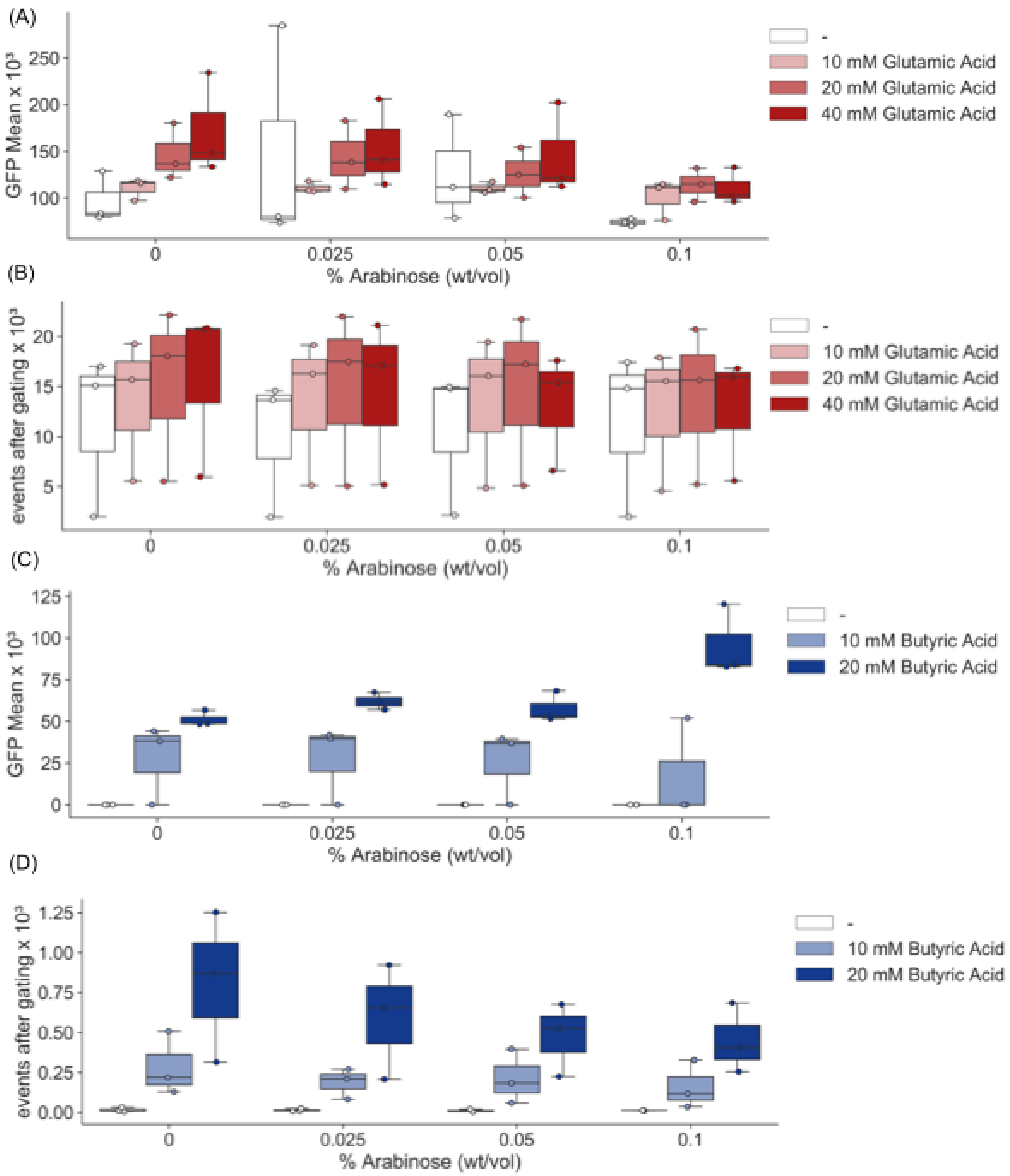


Supplemental Figure 9

***Supplemental Figure 9. Single plasmid system calibration:*** Two-dimensional dose response curves with increasing concentrations of % arabinose, which induces expression of the response regulator, and increasing concentration of the two-component system inducer. (A) Glutamic acid responsive RR, PP_1066, driven by pBAD promoter fluorescence was measured by flow cytometry and measurements are reported as GFP Mean x $10^3$after gating(A) and number of events after gating, if the number of events was < 150, we excluded the data and set the GFP mean to 0 (C). Butyric acid responsive RR, (B) PP_3551, driven by pBAD promoter driven by pBAD promoter fluorescence was measured by flow cytometry and measurements are reported as GFP Mean x $10^3$after gating (C) and number of events after gating, if the number of events was < 150, we excluded the data and set the GFP mean to 0 (D). Center line, median; box limits, upper and lower quartiles; whiskers, 1.5x interquartile range; points with black diamonds, outliers; n = 3.

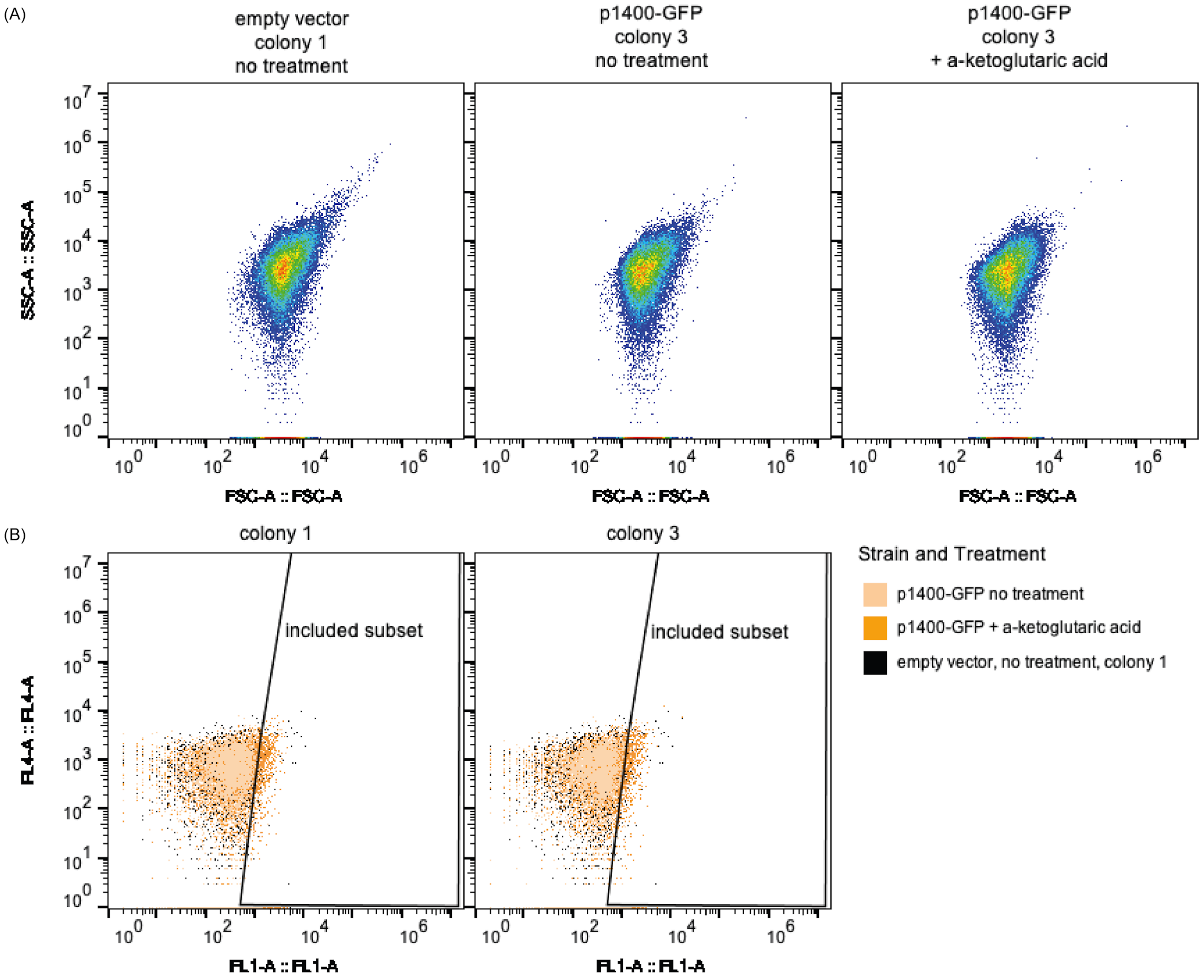


***Supplemental Figure 10. Flow cytometry gating strategy***: (A) All samples exhibited similar FSC-SSC profiles as compared to the empty vector negative control. (B) Assuming the empty vector fluorescence profile designates *P. putida* KT2440's native background fluorescence levels, we gated all samples and removed all events with fluorescence values within the empty vector's distribution. To do so, we plotted the RFP channel (y-axis) against the GFP channel (x-axis) to find the optimal gate for event inclusion. The gating strategy is demonstrated with two biological replicates bearing a-ketoglutaric acid-activated GFP reporter plasmid, p1400-GFP, with or without treatment. The included subset represents a shift in the overall distribution profile of GFP expression.

| Database | unique taxa in rank below | max unique species per rank below | Figure |
|---|---|---|---|
| **ActinobacteriaDB** | 6 | 7 | SF3B |
| **ActinobacteriaDB_2** | 6 | 7 | SF3A |
| **AlphaproteobacteriaDB_1** | 12 | 2 | Figure 2 |
| **AlphaproteobacteriaDB_2** | 12 | 5 | SF3C |
| **BacteriaDB_1** | 35 | 9 | Figure 1D, SF1A, SF2A,B |
| **BacteriaDB_2** | 34 | 7 | SF1 C |
| **BacteriaDB_3** | 34 | 7 | SF1 B |
| **BacteroidetesDB** | 6 | 2 | SF3B |
| **BacteroidetesDB_2** | 6 | 2 | SF3A |
| **BetaproteobacteriaDB_1** | 5 | 2 | Figure 2 |
| **BetaproteobacteriaDB_2** | 5 | 4 | SF3C |
| **ChloroflexiDB** | 8 | 2 | SF3B |
| **ChloroflexiDB_2** | 8 | 2 | SF3A |
| **DeltaproteobacteriaDB_1** | 7 | 4 | Figure 2 |
| **DeltaproteobacteriaDB_2** | 7 | 4 | SF3C |
| **FirmicutesDB** | 6 | 4 | SF3B |
| **FirmicutesDB_2** | 6 | 4 | SF3A |
| **GammaproteobacteriaDB_1** | 19 | 3 | Figure 2 |
| **GammaproteobacteriaDB_2** | 19 | 4 | SF3C |
| **ProteobacteriaDB_1** | 9 | 5 | Figure 1D, SF1A, SF2A,B |
| **ProteobacteriaDB_2** | 9 | 9 | SF1 C |
| **ProteobacteriaDB_3** | 9 | 10 | SF1 B |
| **PseudomonasDB_1** | 51 | 1 | Figure 1D, SF1A, SF2A,B |
| **PseudomonasDB_2** | 51 | 1 | SF1 C |
| **PseudomonasDB_3** | 51 | 1 | SF1 B |

**Table S1.**
***Summary of REC domain random sampling.*** Databases were created by randomly sampling REC domains from curated genomes found in the microbial signal transduction database. To control for overrepresentation of subclades due to more representative species with full genome sequences in any given subclade, species were sampled evenly from the taxonomic rank below the one represented in the plot title (e.g. The taxonomic rank below kingdom is phylum; the maximum of species from each bacterial phylum were randomly sampled to generate the bacteria dataset). This table summarizes the number of taxa that are represented in the REC sequence landscapes in each indicated figure using each indicated database and the maximum number of species per rank below used to generate the data. The perplexity, or expected size of the clusters in the dimensional reduction of the REC sequence alignments, is defined by the number of "unique taxa below rank" for a given database.

| RR Locus Tag | Locus Tag of Genes predicted to be regulated by RR based on DAP-seq data |
|---|---|
| AO356_18205 | AO356_18230, AO356_00230 |
| PP_1066 | PP_1071, PP_2453 |
| Pf6N2E2_5565 | Pf6N2E2_5570 |
| Psest_4267 | Psest_4268, Psest_3930 |
| PP_0263 | PP_1188 |
| AO356_10065 | AO356_18980 |
| Pf6N2E2_5487 | Pf6N2E2_5486;Pf6N2E2_5487 |
| PP_1401 | PP_1400, PP_2426, PP_2790, PP_2909, PP_5045 |
| Psest_4132 | Psest_0085, Psest_0404, Psest_1086;Psest_1087, Psest_1255, Psest_3116, Psest_3304;Psest_3305, Psest_3379,Psest_4201 |
| AO356_17800 | AO356_01395, AO356_08640;AO356_08645, AO356_09700, AO356_09855, AO356_16320;AO356_16325, AO356_17010, AO356_17245, AO356_17790, AO356_19350 AO356_18980, AO356_19350, AO356_20245, AO356_23670, AO356_25920, AO356_30230 |

**Table S2.**
Summarized results of DAP-seq experiments showing predicted targets of RRs referenced in this study.

| Strain | Minimal Media Name | Recipe | Reference (doi) |
|---|---|---|---|
| *P. stutzeri RCH2* | UGA | 4.7 mM ammonium chloride, 1.3 mM potassium chloride, 2 mM magnesium sulfate, 0.1 mM calcium chloride, 0.3 mM sodium chloride, 5 mM sodium dihydrogen phosphate, 20 mM sodium lactate, 25 mM MOPS. Vitamins and minerals were added as described by Widdel and Bak (doi: 10.1007/978-1-4757-2191-1_21) | 10.1128/AEM.01845-16, 10.1007/978-1-4757-2191-1_21 |
| *P. putida KT2440* | M9 | 1 g/L Ammonium sulfate, 1.5 g/L Potassium phospahte monobasic, 3.54 g/L Sodium phospahte dibasic, 0.2 g/L Magnesium sulfate heptahydrate, 0.01 g/L Calcium chloride, 0.06 g/L ammonium ferric citrate, and trace elements (0.3 mg/mL H3BO3, 0.2 mg/mL CoCl2·6H2O, 0.1 mg/L ZnSO4·7H2O, 0.03 mg/LMnCl2·4H2O, 0.03 mg/LNaMoO4·2H2O, 0.02 mg/L NiCl2·6H2O , 0.01 mg/L CuSO4·5H2O ). | 10.1007/s11274-007-9480-x 10.1002/mabi.200600187 |

**Table S3.**
Minimal media recipes used in this study.

**Table S4. (separate file)**
List of primers used in this study.

**Supplementary Methods**

***Buffer Recipes***

All buffers are made in Tris Buffer Saline pH 7.4 (1X TBS), however, TBS can be substituted with another buffering solution, such as HEPES or phosphate buffer saline.

| **Buffer Name** | **Recipe** |
|---|---|
| Lysis Buffer | 100 µM PMSF (Millipore Sigma, Burlington MA), 2.5 units/mL Benzonase nuclease (Millipore Sigma, Burlington MA), 1 mg/mL Lysozyme (Millipore Sigma, Burlington MA), in 1X TBS |
| Wash Buffer | 10 mM Imidazole, 0.1% Tween 20 in 1X TBS |
| DNA Binding Buffer | 10 mM magnesium chloride, 0.4 ng/µL AL-DNA, with or without 50 mM acetyl phosphate in 1X TBS |
| Enrichment Buffer | 10 mM Imidazole, 0.1% Tween 20 in 1X TBS |
| Elution Buffer | 180 mM Imidazole in 1X TBS |
| Storage Buffer | 20 % (v/v) ethanol |
| Recharge Equilibration Buffer | 1X TBS |
| Recharge Stripping Buffer | 50 mM ethylenediaminetetraacetic acid (EDTA) and 500 mM NaCl in 2X TBS |
| Recharge Salt Buffer | 500 mM sodium chloride in 2X TBS |
| Recharge Solution | 100 mM Cobalt chloride in water |

**Step by step methods for automated DNA affinity purification**

*DNA preparation for NGS*

1. Bacterial strains can be cultured in either LB or minimal media (see supplementary table 2)
2. Purify genomic DNA (by any available method). *Note: DNA should be stored in a buffer containing EDTA to ensure stability during shearing.*
3. Shear the genomic DNA with covaris miniTUBE (Covaris, Woburn, MA) to an average of 200 bp.
4. *Quality Control (QC) Step*: Check DNA quality by Bioanalyzer high sensitivity DNA kit (Agilent, Santa Clara)
5. Adapter ligate 1µg sheared DNA with NEBnext Ultra ii Library Preparation kit (New England Biolabs, Ipswich, MA) or other library preparation kit with input range > 0.5 µg. *Note: STOP before PCR step.*
6. *QC*: Check DNA quality by Bioanalyzer high sensitivity DNA kit (Agilent, Santa Clara)
7. DNA can be stored at -20˚C.

*RR-expression*

1. Use standard cloning techniques to build RR expression strains (N-terminal 6xHis-RR in Pet28), and transform plasmids into a heterologous expression host, such as BL21 de3.

2. Grow strains overnight (ON) in LB in a 96 deep-well plate. Back-dilute 5 µL of ON culture into wells of 995 µL autoinduction media (Zyp-5052). Grow strains at 37˚C shaking at 250 RPM for 5-6 hours, then transfer plates to 17˚C, 250 RPM for overnight growth.
3. Pellet cultures in a centrifuge at 3214 x g, pour off the supernatant and store at -20˚C for no more than 1 week.

*Automated pipeline for DNA affinity purification*

An automated pipeline for DAP was built on Beckman Coulter Biomek FX, using Phynexus 5 µL IMAC resin tips recharged with cobalt chloride (see step 8) to decrease the required concentration of imidazole and for improved protein purity.

1. Lysis
    a. Lyse 1 mL cell pellets with 300 µL **Lysis Buffer** for 1 hour at 30˚C.
    b. Clarify lysates by centrifugation at 3214 x g. (*Note: the centrifuge can be set to 4˚C if protein degradation is a concern*).
    c. To ensure there is no remaining cell debris, further clarify the lysate using 20 µm pore 96-well filter plates by centrifugation at 1800 x g (*Note: the centrifuge can be set to 4˚C if protein degradation is a concern*).
2. Equilibration:
    a. **Storage Buffer** is expelled from IMAC resin tips by performing a high pressure blow-out over the **Storage Buffer** reservoir.
    b. IMAC resin tips are washed or remaining ethanol by suspending and dispensing 100 µL water for 1 minute.
    c. IMAC resin tips are equilibrated by suspending and dispensing 100 µL **Wash Buffer** for 1 minute.
3. Protein Binding:
    a. IMAC resin tips bind metal affinity tagged protein by suspending and dispensing 100 µL clarified lysates (stored in 96-well plate from step 1a-c) for 10 minutes.
    b. The tips are then washed by suspending and dispensing 100 µL **Wash Buffer** washed for one minute in the wells of a 96-well plate containing 300 µL **Wash Buffer** per well.
    c. *Note: to avoid cross-contamination do not use a reservoir for any wash or enrichment steps.*
4. DNA Binding:
    a. Protein bound to IMAC resin binds target DNA by suspending and dispensing 50µL **DNA Binding Buffer** in the wells of a 96-well plate containing 60 µL **Wash Buffer** per well.
    b. Bound DNA is then enriched by suspending and dispensing 100 µL **Enrichment Buffer** in the wells of 3 separate 96-well plates containing 300 µL **Enrichment Buffer** per well for 5 minutes per plate.
5. Tip Drying:
    a. To ensure no residual buffers from the previous steps remain on the IMAC resin tips before elution, tips are dried with a custom protocol, in which liquid remaining in the tip undergoes a high pressure blow-out.
    b. The tips are then touched to a kim-wipe stabilized with a tip-box lid. The kim-wipe collects any residual liquid and the tips are dried for the next steps.
6. Elution:
    a. Protein-DNA complex bound to IMAC resin is eluted by suspending and dispensing 25µL µL **Elution Buffer** in the wells of a 96-well plate containing 25 µL **Elution Buffer** per well.
    b. Eluted Protein-DNA can be stored at -20˚C for 1 week before library preparation.
    c. *Optional QC*: Sample a few wells to check for protein expression by western blot
    d. *Note: Downstream removal of salts and imidazole are not necessary at this stage, as this protocol leverages a reagent tolerant PCR master-mix for library preparation.*
7. Final washing and storage:
    a. IMAC resin tips are then washed in a reservoir containing 150 mL **Elution Buffer,** followed by a reservoir containing 150 mL water.
    b. The tips are then washed in a reservoir with 150 mL **Storage Buffe**r.

   c. Before completing the wash, the tips suspend 150 µL **Storage Buffer** without dispensing and return to their original box.
   d. The tips are then wrapped in parafilm and stored at 4˚C. Tips can be recharged and reused up to 10 times (see step 8).
8. Recharging IMAC resin tips:
   a. Equilibrate tips by suspending and dispensing 100 µL **Recharge Equilibration Buffer** stored in 50 mL reservoir for 1 minute.
   b. Suspend and dispense 100 µL **Recharge Stripping Buffer** stored in a 50 mL reservoir for 1 minute.
   *c.* *Optional:* Suspend and dispense 100 µL 1 N NaOH stored in a 50 mL reservoir for 1 minute. Equilibrate tips again by suspending and dispensing 100 µL **Recharge Equilibration Buffer** stored in 50 mL reservoir for 1 minute.
   d. *Optional:* Suspend and dispense 100 µL **Recharge Salt Buffer** stored in a 50 mL reservoir for 1 minute. Equilibrate tips again by suspending and dispensing 100 µL **Recharge Equilibration Buffer** stored in 50 mL reservoir for 1 minute.
   e. Suspend and dispense 100 µL **Recharge Equilibration Buffer** stored in a 50 mL reservoir for 1 minute.
   f. Suspend and dispense 100 µL **Recharge Solution** stored in a 50 mL reservoir for 1 minute.
   g. Suspend and dispense 100 µL **Recharge Equilibration Buffer** stored in a 50 mL reservoir for 1 minute.
   h. The tips are then washed in a reservoir with 150 mL **Storage Buffe**r.
   i. Before completing the wash, the tips suspend 150 µL **Storage Buffer** without dispensing and return to their original box.
   j. The tips are then wrapped in parafilm and stored at 4˚C. Tips can be recharged and reused up to 10 times.
